# Two-component spike nanoparticle vaccine protects macaques from SARS-CoV-2 infection

**DOI:** 10.1101/2020.11.07.365726

**Authors:** Philip J. M. Brouwer, Mitch Brinkkemper, Pauline Maisonnasse, Nathalie Dereuddre-Bosquet, Marloes Grobben, Mathieu Claireaux, Marlon de Gast, Romain Marlin, Virginie Chesnais, Ségolène Diry, Joel D. Allen, Yasunori Watanabe, Julia M. Giezen, Gius Kerster, Hannah L. Turner, Karlijn van der Straten, Cynthia A. van der Linden, Yoann Aldon, Thibaut Naninck, Ilja Bontjer, Judith A. Burger, Meliawati Poniman, Anna Z. Mykytyn, Nisreen M. A. Okba, Edith E. Schermer, Marielle J. van Breemen, Rashmi Ravichandran, Tom G. Caniels, Jelle van Schooten, Nidhal Kahlaoui, Vanessa Contreras, Julien Lemaître, Catherine Chapon, Raphaël Ho Tsong Fang, Julien Villaudy, Kwinten Sliepen, Yme U. van der Velden, Bart L. Haagmans, Godelieve J. de Bree, Eric Ginoux, Andrew B. Ward, Max Crispin, Neil P. King, Sylvie van der Werf, Marit J. van Gils, Roger Le Grand, Rogier W. Sanders

## Abstract

The SARS-CoV-2 pandemic is continuing to disrupt personal lives, global healthcare systems and economies. Hence, there is an urgent need for a vaccine that prevents viral infection, transmission and disease. Here, we present a two-component protein-based nanoparticle vaccine that displays multiple copies of the SARS-CoV-2 spike protein. Immunization studies show that this vaccine induces potent neutralizing antibody responses in mice, rabbits and cynomolgus macaques. The vaccine-induced immunity protected macaques against a high dose challenge, resulting in strongly reduced viral infection and replication in upper and lower airways. These nanoparticles are a promising vaccine candidate to curtail the SARS-CoV-2 pandemic.

## Introduction

Severe acute respiratory syndrome coronavirus 2 (SARS-CoV-2) has rapidly spread across the globe and infected more than 46 million individuals since late 2019 (https://covid19.who.int/). SARS-CoV-2 causes coronavirus disease 2019 (COVID-19), which manifests itself as a mild respiratory illness in most infected individuals, but can lead to acute respiratory distress syndrome and death in a significant percentage of cases. As of 1 October 2020, COVID-19 has caused over 1 million casualties and continues to place a significant burden on healthcare systems and economies worldwide. Hence, the development of a safe and effective vaccine that can prevent SARS-CoV-2 infection and transmission has rapidly become top priority.

Recent studies suggest that SARS-CoV-2-specific neutralizing antibody (NAb) titers are an important immune correlate of protection (Addetia et al., 2020; Yu et al., 2020). This is supported by several passive immunization studies showing that administration of potent neutralizing SARS-CoV-2-specific monoclonal antibodies (mAbs) can significantly reduce lung viral loads (Cao et al., 2020; Rogers et al., 2020). Thus, a successful vaccine will likely need to induce a potent NAb response. The main target for NAbs on SARS-CoV-2 is the spike (S) protein. This homotrimeric glycoprotein is anchored in the viral membrane and consists of the S1 domain, containing the receptor-binding domain (RBD) for the host cell receptor angiotensin converting enzyme-2 (ACE2), and the S2 domain, which contains the fusion peptide. Upon binding to ACE2, the prefusion S protein undergoes several structural changes that induce a shift to a postfusion state that enables merging of the viral envelope and host cell membrane (Shang et al., 2020). As most NAb epitopes are presented on the prefusion conformation, vaccine candidates should include the prefusion S protein, which as for other class I viral fusion proteins (Krarup et al., 2015; Sanders et al., 2002), can be stabilized by appropriately positioned proline substitutions (Pallesen et al., 2017; Walls et al., 2020a; Wrapp et al., 2020).

More than 200 candidate vaccines are currently under preclinical or clinical evaluation (https://www.who.int/publications/m/item/draft-landscape-of-covid-19-candidate-vaccines). Recombinant subunit vaccines provide a welcome alternative to the inactivated, viral vector and RNA-based vaccines that are currently in Phase III clinical trials, as they have a track record of safety and efficacy. In addition, recombinantly expressed S proteins represent the most efficient antigens to induce potent NAb responses, as recently reported in non-human primate studies (Guebre-Xabier et al., 2020; Liang et al., 2020; Tian et al. 2020).

A well established strategy to generate strong humoral immune responses against soluble antigens is multivalent antigen display (Bachmann and Jennings, 2010; Moyer et al., 2016). Presenting a high density of antigen on a repetitive array facilitates numerous immunological processes such as B cell activation, lymph node trafficking and retention on follicular dendritic cells (Kelly et al., 2019; Tokatlian et al., 2019). Among the several nanoparticle designs that are currently being employed to present viral glycoproteins, self-assembling protein nanoparticle systems represent promising platforms as they allow for efficient and scalable nanoparticle assembly (López-Sagaseta et al., 2016). Homomeric protein complexes such as the 24-subunit ferritin and 60-subunit mi3 nanoparticles self assemble intracellularly and have been used to display immunogens such as influenza HA, HIV-1 Env, malaria CyRPA and more recently SARS-CoV-2 S (Bruun et al., 2018; He et al., 2020; Kanekiyo et al., 2013; Powell et al., 2020; Sliepen et al., 2015). Recently, two-component 120 subunit icosahedral nanoparticles, such as the I53-50 and dn5 designs, have been developed that self-assemble *in vitro,* allowing for controlled nanoparticle formation. We and others have been using these nanoparticles to multivalently present trimeric type 1 viral fusion proteins of HIV-1, respiratory syncytial virus (RSV) and influenza (Boyoglu-Barnum et al., 2020; Brouwer et al., 2019;Marcandalli et al., 2019). Presentation of these immunogens on two-component protein nanoparticles significantly improved NAb titers, which supports pursuing this platform for the generation of nanoparticle immunogens displaying prefusion SARS-CoV-2 S proteins.

Here, we describe the generation and characterization of two-component protein nanoparticles displaying stabilized prefusion SARS-CoV-2 S proteins. Immunization studies in mice, rabbits and macaques demonstrated that these nanoparticles induce robust humoral responses. Vaccinated macaques challenged with a high dose of SARS-CoV-2 virus had strongly reduced viral loads in both the upper and lower respiratory tracts, and were preserved from lung lesions when compared to unvaccinated animals.

## Results

### SARS-CoV-2 S proteins can be displayed on two component I53-50 nanoparticles

The computationally designed I53-50 nanoparticle (I53-50NP) constitutes twenty trimeric (I53-50A or variants thereof) and twelve pentameric (I53-50B.4PT1) subunits which self-assemble to form monodisperse icosahedral particles with a diameter of approximately 30 nm (Bale et al., 2016). To generate I53-50NPs presenting SARS-CoV-2 S proteins, we genetically fused the C-terminus of the previously described stabilized prefusion S protein to the N-terminus of the I53-50A variant, I53-50A.1NT1, using a glycine-serine linker (Figure 1A) (Brouwer et al.,2020). The fusion protein was purified from transfected human embryonic kidney (HEK) 293F cell supernatant using Ni-NTA purification followed by size-exclusion chromatography (SEC). Collection of the appropriate SEC fractions yielded ~2 mg/L of trimeric SARS-CoV-2 S-I53-50A.1NT1 fusion protein (Figure 1B). To initiate assembly of nanoparticles presenting SARS-CoV-2 S proteins (SARS-CoV-2 S-I53-50NPs), the pooled trimer fractions were incubated overnight at 4°C with I53-50B.4PT1 in an equimolar ratio. The nanoparticle preparation was further purified using an additional SEC step to remove unassembled components. Negative-stain electron microscopy (nsEM) of the pooled higher molecular weight fractions revealed a considerable portion of monodisperse and well-ordered icosahedral nanoparticles (Figure 1C). Bio-layer-interferometry (BLI)-based binding experiments with a panel of SARS-CoV-2 S protein-specific monoclonal NAbs, previously isolated from recovered COVID-19 patients (Brouwer et al., 2020), showed strong binding of RBD-specific COVA1-18, COVA2-02, COVA2-15 and COVA2-39, and N-terminal domain (NTD)-specific COVA1-22, to trimeric SARS-CoV-2 S-I53-50A.1NT1 and SARS-CoV-2 S-I53-50NP (Figure 1D). This suggests that presentation of SARS-CoV-2 S on I53-50NPs did not compromise the structure of these S protein epitopes. Altogether, SEC, nsEM and BLI confirmed the successful generation of nanoparticles presenting multiple copies of the SARS-CoV-2 S protein.

**Figure 1.**
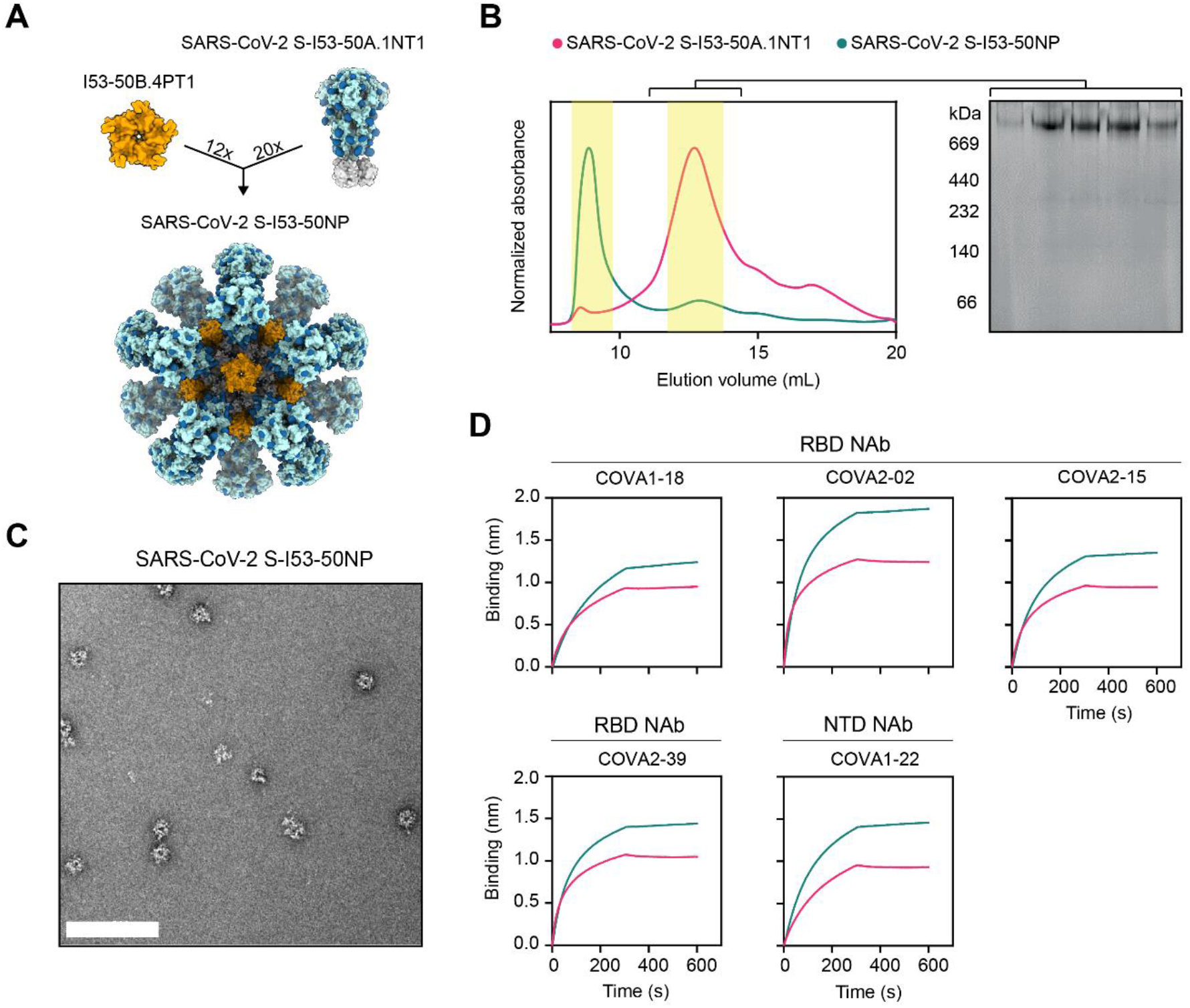
Biophysical and antigenic characterization of SARS-CoV-2 S-I53-50NPs. (A) Schematic representation of 20 copies of SARS-CoV-2 S-I53-50A.1NT1 (SARS-CoV-2 S in light blue with glycans in dark blue, I53-50A.1NT1 in white) and 12 copies of I53-50B.4PT1 assembling into SARS-CoV-2 S-I53-50NP. (B) Size exclusion chromatograms of SARS-CoV-2 S-I53-50A.1NT1 (magenta) and SARS-CoV-2 S-I53-50NP (green) run over a Superose 6 increase 10/300 GL column. The yellow columns specify the SEC fractions that were collected, pooled and used. Blue native gel of pooled SARS-CoV-2 S-I53-50A.1NT1 SEC fractions. (C) Negative-stain EM analysis of assembled SARS-CoV-2 S-I53-50NPs. The white bar represents 200 nm. (D) BLI sensorgrams showing the binding of multiple SARS-CoV-2 NAbs to SARS-CoV-2 S-I53-50A.1NT1 (magenta) and SARS-CoV-2 S-I53-50NP (green).

As approximately a third of the mass of the SARS-CoV-2 S protein consists of N-linked glycans, we sought to identify the site-specific glycosylation of S protein presented on I53-50NPs using liquid chromatography-mass spectrometry (LC-MS). All sites presented high levels of occupancy, with only N1074 displaying 15% of sites lacking an N-linked glycan (Figure S1A). The compositions of glycans present at the 19 N-linked glycan sites on the S protein were determined and revealed a diverse range of structures (Figure S1B). Underprocessed oligomannose-type glycans were observed at sites N61, N234, N717, N801 and to a lesser extent at N1098. The average oligomannose-type glycan content across all sites was 11%. Processed complex-type glycans were observed at all sites and were highly fucosylated (73%), but mostly lacked sialylation (8%) (Figures S1A and S1B). The glycoforms present on SARS-CoV-2 S-I53-50NPs are more processed compared to other recombinant S protein immunogens (Watanabe et al., 2020), but are reminiscent of the composition on S proteins presented on virions (Yao et al., 2020).

### SARS-CoV-2 S-I53-50NPs enhance activation of SARS-CoV-2 S protein-specific B cells *in vitro*

Multivalent display of antigens can enhance cognate B cell activation over soluble antigen (Boyoglu-Barnum et al., 2020; Kanekiyo et al., 2013; Marcandalli et al., 2019; Veneziano et al., 2020). To assess if the same would apply for SARS-CoV-2 S-I53-50NPs, we generated B cells that expressed B cell receptors (BCRs) for previously described RBD-targeting monoclonal NAbs and measured their activation by monitoring Ca^2+^ influx *in vitro* (Brouwer et al., 2020). Soluble trimers only weakly activated COVA1-18 expressing B cells at the highest concentration used (5 μg/mL SARS-CoV-2 S-I53-50A.1NT1), while an equimolar amount of SARS-CoV-2 S presented on I53-50NPs efficiently activated the same B cells (Figure 2). COVA2-15-expressing B cells were activated by soluble SARS-CoV-2 S-I53-50A.1NT1 trimers, but markedly more efficiently so by SARS-CoV-2 S-I53-50NP. The more efficient activation of COVA2-15-expressing B cells, than those expressing COVA1-18, may be explained by the fact that COVA2-15 can interact with the RBD in both the up and down state, which may result in higher avidity interactions (Brouwer et al., 2020).In control experiments, I53-50NPs displaying soluble HIV-1 envelope glycoproteins (BG505 SOSIP) did not activate any of the B cell lines. We conclude that SARS-CoV-2 S-I53-50NPs considerably improve the activation of SARS-CoV-2-specific B cells compared to soluble trimers.

**Figure 2.**
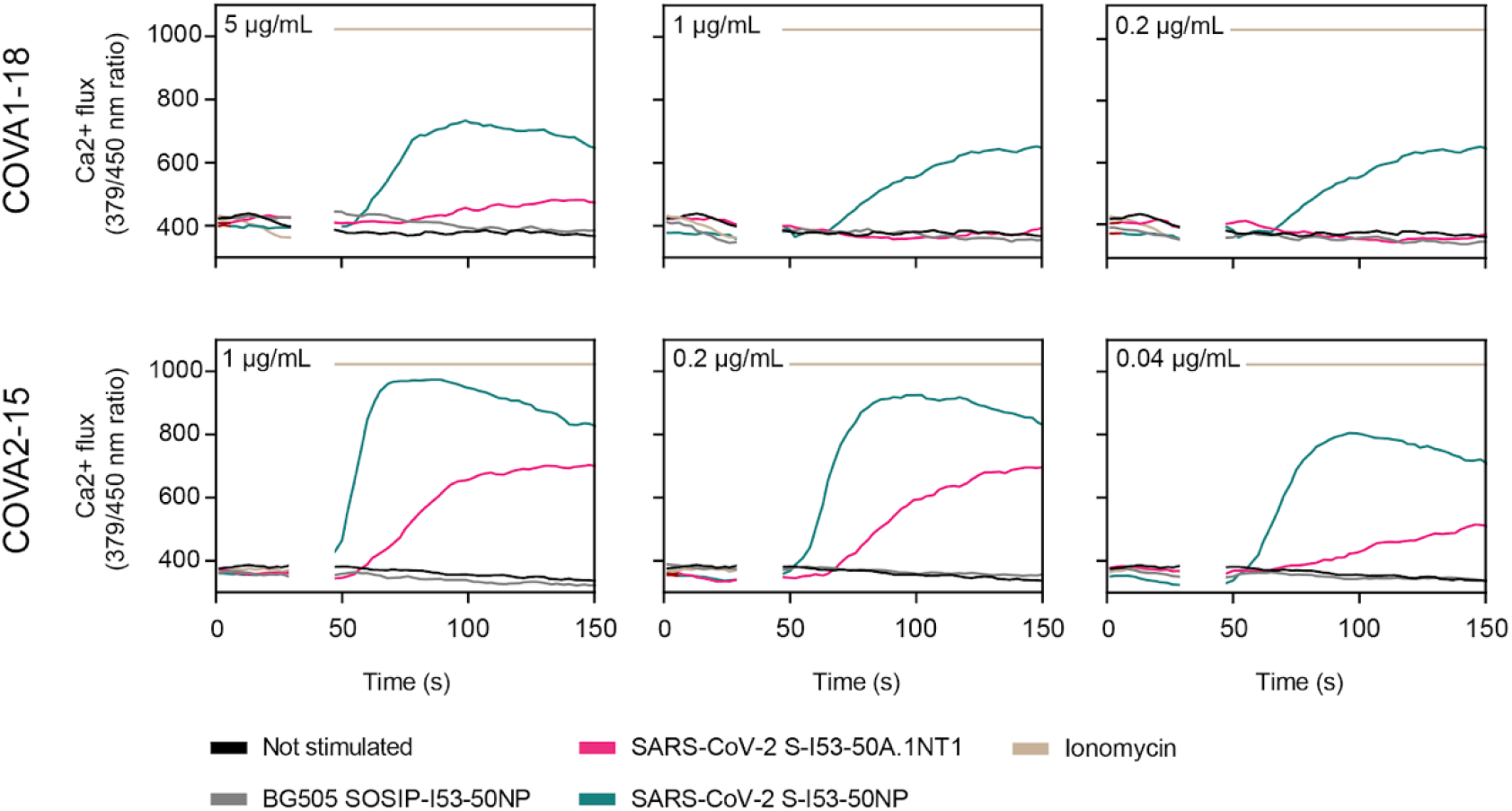
*In vitro* B cell activation by SARS-CoV-2 S-I53-50A.1NT1 and SARS-CoV-2 S-I53-50NPs. B cells expressing the SARS-CoV-2 S protein-specific NAbs COVA1-18 (top) or COVA2-15 (bottom) as BCRs were incubated with either SARS-CoV-2 S-I53-50A.1NT1 (magenta), SARS-CoV-2 S-I53-50NP (green), lonomycin (beige), or BG505 I53-50NP (grey), or not stimulated (black). The experiments were performed with 5, 1, 0.2 or 0.04 μg/mL of SARS-CoV-2 S-I53-50A.1NT1, as indicated in the top left corner of each graph, or the equimolar amount of SARS-CoV-2 S or BG505 SOSIP on I53-50NPs. Ionomycin was used at 1 mg/mL as a positive control.

### SARS-CoV-2 S-I53-50NP vaccination induces robust NAb responses in small animal models

We assessed the immunogenicity of SARS-CoV-2 S-I53-50NPs in two small animal models. Eight BALB/c mice were immunized with 10 μg of SARS-CoV-2 S presented on I53-50NPs, adjuvanted in polyinosinic-polycytidylic acid (Poly-IC). In addition, five New Zealand White rabbits were immunized with 30 μg of SARS-CoV-2 S presented on I53-50NPs, adjuvanted in squalene emulsion. Mice and rabbits received three subcutaneous and intramuscular immunizations, respectively, at week 0, 4 and 12 (Figure 3A).

**Figure 3.**
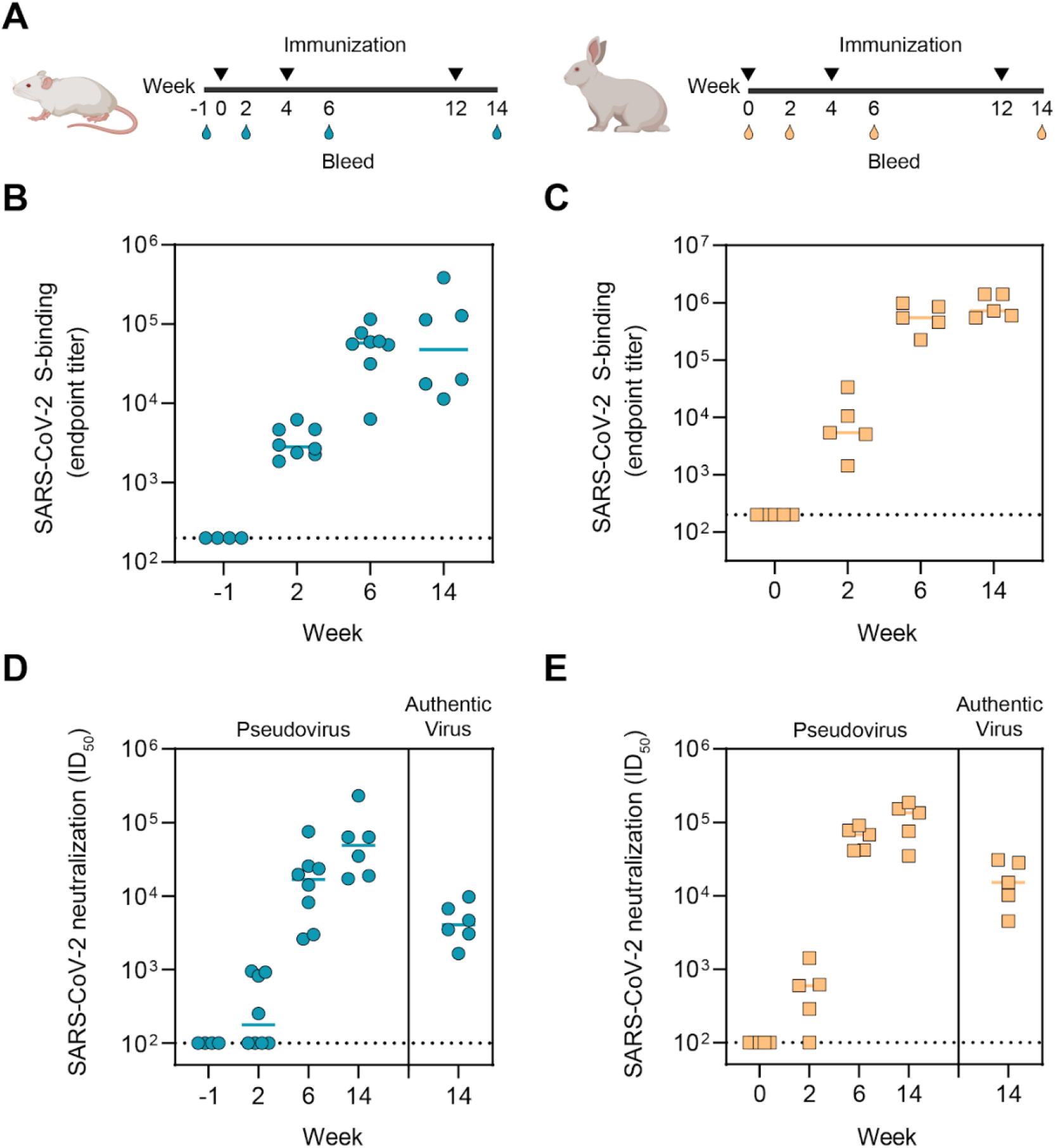
Immunogenicity of SARS-CoV-2 S-I53-50NPs in mice and rabbits. (A) Study schedule in mice (left) and rabbits (right). Black triangles indicate the timepoints of immunization and drops indicate the bleeds. (B) ELISA endpoint titers for SARS-CoV-2 S protein-specific IgG in mice. Due to limited volumes of sera at week-1, random pairs of mice sera were pooled. (C) ELISA endpoint titers for SARS-CoV-2 S protein-specific IgG in rabbits. (D) SARS-CoV-2 pseudovirus and authentic virus neutralization titers in mice. (E) SARS-CoV-2 pseudovirus and authentic virus neutralization in rabbits. (B, C) Week-1 samples were pooled to make 4 samples. At week 6, 2 animals were sacrificed. (B-E) The median titers are indicated by a bar.

Two weeks after the first immunization, mice and rabbits induced detectable SARS-CoV-2 S protein-specific immunoglobulin G (IgG) titers, as determined by an enzyme-linked immunosorbent assay (ELISA), with a median endpoint binding titer of 2,920 and 5,105 respectively. In mice, median endpoint titers were further boosted after the second immunization to 57,943 at week 6 and slightly decreased after the third, to 47,643 at week 14 (Figure 3B). In rabbits, median endpoint titers were boosted to 544,503 at week 6 and 594,292 at week 14 (Figure 3C). Neutralization of SARS-CoV-2 pseudovirus was already detectable in the majority of mice and rabbit sera two weeks after the first immunization. NAb titers, which are represented by the inhibitory dilutions at which 50% neutralization is attained (ID_50_ values), increased to a median of 16,792 at week 6 and 49,039 at week 14 in mice (Figure 3D and table S1). In rabbits, NAb titers were boosted to a median ID_50_ of 68,298 and 135,128 at week 6 and 14, respectively (Figure 3E and Table S1). Neutralization titers of authentic SARS-CoV-2 virus at week 14 reached a median ID_50_ of 4,065 and 15,110 in mice and rabbits, respectively (Figures 3D and 3E, and Table S1). Collectively, we showed that SARS-CoV-2 S-I53-50NPs were able to induce robust binding and NAb responses in both mice and rabbits after two and three immunizations.

### SARS-CoV-2 S-I53-50NP vaccination induces potent humoral responses in cynomolgus macaques

The high binding and neutralization titers in mice and rabbits supported subsequent assessment of the immunogenicity of SARS-CoV-2 S-I53-50NPs in cynomolgus macaques; an animal model that is immunologically closer to humans. Six cynomolgus macaques were immunized with 50 μg of SARS-CoV-2 S-I53-50NPs formulated in monophospholipid A (MPLA) liposomes by the intramuscular route at week 0, 4 and 10 (Figure 4A).

**Figure 4.**
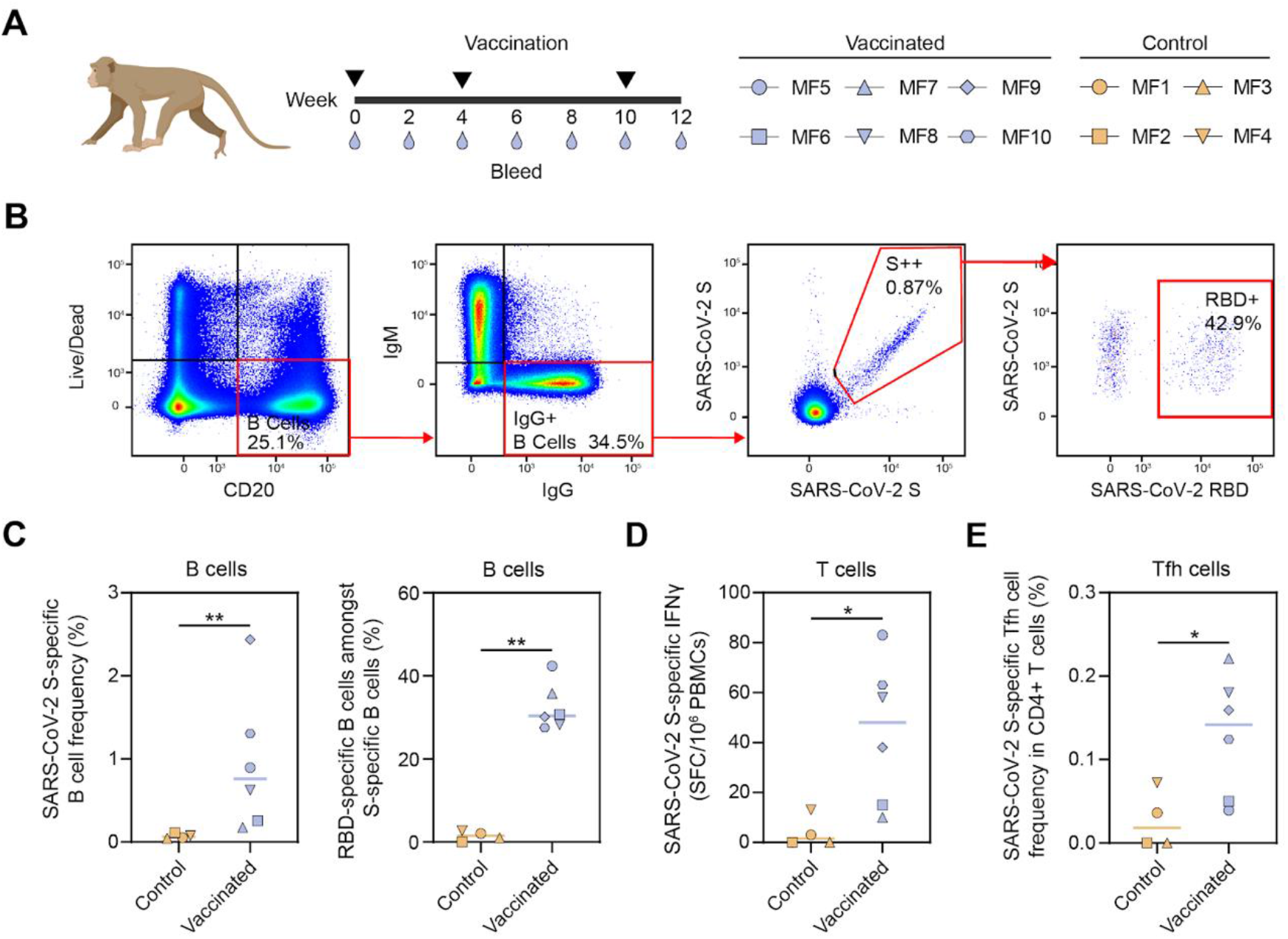
SARS-CoV-2 S protein-specific B and T cell responses induced by SARS-CoV-2 S-I53-50NPs in cynomolgus macaques. (A) Vaccination schedule in cynomolgus macaques. Black triangles indicate the timepoints of vaccination and drops mark the bleeds. The symbols identifying individual macaques are used consistently throughout figures 4–6. (B) Representative gating strategy, depicting the analysis of SARS-CoV-2 S protein- and RBD-specific IgG+ B cells, shown for one vaccinated macaque. The lymphocyte population was selected and doublets were excluded (not shown). Gating strategy from the left to the right: viable B cells (Live/dead-CD20+), IgG+ B cells (IgM-IgG+), SARS-CoV-2 S protein (double probe staining) and RBD (single probe staining)-specific IgG+ B cells. (C) SARS-CoV-2 S protein-specific B cell frequencies within the IgG+ population, in control and vaccinated macaques (left). Percentages of SARS-CoV-2 RBD-specific B cells within the population of SARS-CoV-2 S protein-specific IgG+ B cells (right). (D) Number of IFNy secreting cells after ex vivo stimulation with SARS-CoV-2 S protein as analysed by ELISpot and plotted as spot forming cells (SFC) per 1.0×10^6^ PBMCs. (E) Frequency of SARS-CoV-2 S protein-specific Tfh cells (CD69+ CD154+ CXCR5+) in the total CD4+ T cell population. PBMCs were stimulated overnight with SARS-CoV-2 S protein and Tfh activation was assessed the next day by analyzing CD69 and CD154 expression by flow cytometry. The gating strategy to identify this population is shown in Figure S2. (C-E) Bars indicate median. Groups were compared using the Mann-Whitney U test (*, *p* < 0.05; **, *p* < 0.01).

To analyze the frequency of S protein and RBD-specific IgG-positive B cells in macaques after vaccination, peripheral blood mononuclear cells at week 12 were gated on the expression of CD20 and IgG and stained with fluorescently labeled prefusion SARS-CoV-2 S protein or RBD (Figure 4B). We observed a clear expansion of SARS-CoV-2 S protein-specific B cells by vaccination, which constituted on average about 1% of total B cells (Figure 4C). Within the population of IgG-positive SARS-CoV-2 S protein-specific B cells, on average around 30% bound to the RBD (Figure 4C). SARS-CoV-2 S protein-specific T cells were also markedly expanded, as determined by an enzyme-linked immune absorbent spot assay (ELISpot) (Figure 4D). Furthermore, we observed pronounced expansion of S protein-specific T follicular helper (Tfh) cells (CD69+ CD154+ CXCR5+) within the CD4+ T cell subset, as determined by the activation induced marker (AIM) assay (Figure 4E and Figure S2).

Sera of the immunized macaques exhibited SARS-CoV-2 S protein-specific binding IgG responses with median endpoint titers of 211, 1,601 and 2,190, at week 2, 6, and 12, respectively (Figure 5A). To compare binding titers to SARS-CoV-2 S protein and RBD between sera from vaccinated macaques and those from convalescent humans from the COSCA cohort *(28),* a different ELISA protocol was used. Specifically, binding responses were compared to a standard curve of species-specific polyclonal IgG so that a semi-quantitative measure of specific IgG concentrations could be obtained. Week 6 and 12 sera elicited markedly higher IgG binding titers to SARS-CoV-2 S protein than serum from convalescent humans (Figure 5B). This was also the case for RBD-specific binding titers (Figure 5C).

**Figure 5.**
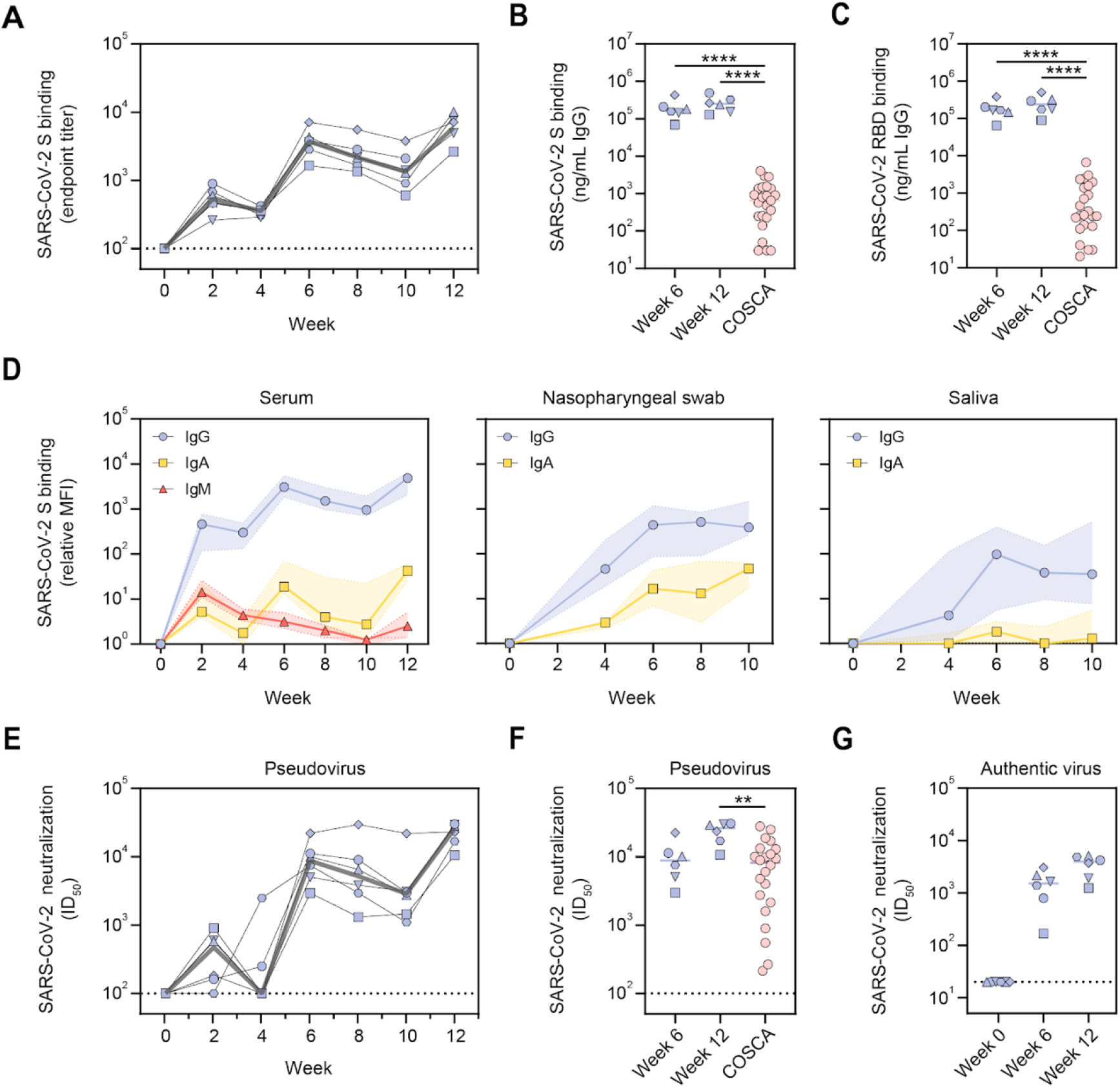
Serological responses induced by SARS-CoV-2 S-I53-50NPs in cynomolgus macaques. (A) ELISA endpoint titers for SARS-CoV-2 S protein-specific IgG. The grey line represents the median titers over time. (B) SARS-CoV-2 S protein-specific binding titers at week 6 and 12 in macaques compared to those in convalescent humans from the COSCA cohort. Patient samples were taken 4 weeks after onset of symptoms. (C) SARS-CoV-2 RBD-specific binding titers at week 6 and 12 in macaques compared to those in convalescent humans from the COSCA cohort. (D) Relative mean fluorescence intensity (MFI) of IgG, IgA and IgM binding to SARS-CoV-2 S protein measured with a Luminex-based serology assay in serum samples, nasopharyngeal swabs and saliva samples. Shown are medians with the shaded areas indicating the interquartile ranges. (E) SARS-CoV-2 pseudovirus neutralization titers. The grey line represents median titers. (F) SARS-CoV-2 pseudovirus neutralization titers at weeks 6 and 12 in macaques compared to those in convalescent humans from the COSCA cohort. (G) SARS-CoV-2 authentic virus neutralization titers at weeks 6 and 12. The bars show the median titers. (B, C, E, G) Groups were compared using the Mann-Whitney U test (**, *p* < 0.01; ****, *p* < 0.0001). The bars indicate median titers. The dotted lines indicate the lowest serum dilution.

Using a custom Luminex bead-based serological assay, we analyzed the induction of several Ig isotypes in serum, nasopharyngeal swab and saliva samples of the vaccinated macaques over time. S-specific IgG levels in serum showed a similar course as observed by ELISA (Figure 5D and Figure S3A). This was also the case for IgA, albeit with a more rapid onset and waning after vaccination. As expected, IgM levels peaked after the first immunization and gradually waned thereafter; presumably a result of Ig class-switching (Figure 5D and Figure S3A). We also observed an increase in S-specific IgG and IgA levels in nasopharyngeal swabs and saliva after consecutive immunizations. This was particularly clear for IgG in nasopharyngeal swabs (Figure 5D and Figure S3B). The results in saliva samples were more variable (Figure 5D and Figure S3C). Thus, in addition to a systemic antibody response, SARS-CoV-2 S-I53-50NPs induced detectable mucosal IgA and IgG responses; a relevant finding considering that the respiratory mucosa is the first port of entry for SARS-CoV-2 (Sungnak et al., 2020). Finally, we analyzed the ability of SARS-CoV-2 S protein-specific serum Abs to bind to immune proteins that play a role in Fc-mediated effector functions. The levels of FcγRIIa, FcγRIIIa and C1q binding tracked with IgG levels, suggesting that the induced IgGs can perform effector functions such as antibody-dependent cellular-cytotoxicity, phagocytosis and complement activation (Figure S3D).

Serum neutralization titers in the vaccinated macaques were overall robust. Already two weeks after the first immunization, macaques induced pseudovirus neutralization with a median ID_50_ of 475. The second immunization increased the neutralization titers to a median ID_50_ of 8,865 which then declined only modestly up to the third immunization. NAb titers were further increased to a median ID_50_ of 26,361 at week 12 (Figure 5E and Table S3). At week 6 the neutralization titers were similar to those in sera from convalescent humans, but neutralization titers at week 12 were significantly higher in vaccinated macaques than convalescent humans (median ID_50_ of 26,361 vs 8,226, *p*= 0.0012) (Figure 5F). Neutralization of authentic SARS-CoV-2 was lower than that of pseudovirus but remained potent (median ID_50_ of 1,501 and 3,942 at week 6 and 12, respectively) (Figure 5G and Table S3).

### SARS-CoV-2 S-I53-50NP vaccination protects cynomolgus macaques from high-dose SARS-CoV-2 challenge

To assess the protective potential of SARS-CoV-2 S-I53-50NPs, vaccinated macaques and contemporaneous control macaques (n=4) were infected with a total dose of 1 x 10^6^ plaqueforming units (PFU) of a primary SARS-CoV-2 isolate (BetaCoV/France/IDF/0372/2020; passaged twice in VeroE6 cells) by combined intra-nasal and intra-tracheal routes at week 12, two weeks after the final immunization. This represents a high dose challenge in comparison with other studies where 10- to 100-fold lower doses were used (van Doremalen et al., 2020;Feng et al., 2020; Guebre-Xabier et al., 2020; Mercado et al., 2020; Patel et al., 2020; Yu et al., 2020). Control animals had high viral load levels in nasopharyngeal and tracheal samples (swabs), as assessed by RT-qPCR for viral RNA, as early as 1 day post exposure (dpe). In tracheal samples, viral loads peaked between 1 and 3 dpe, with a median value of 6.9 log_10_ copies/mL. Subsequently, viral loads progressively decreased and all animals had undetectable viral loads by 14 dpe (Figure 6A and Figure S4). Similar profiles were observed in nasal swabs, although viral loads remained detectable in some macaques at 14 dpe (Figure 6A and Figure S4). Viral loads were markedly lower in rectal samples but stayed above the limit of detection during the course of sampling for 2/4 control macaques (Figure S4A). Viral subgenomic (sg)RNA, which is believed to reflect viral replication, peaked at 2 dpe in tracheal and nasopharyngeal swabs (median viral sgRNA load 5.8 and 7.1 log_10_ respectively) and were still detectable (> 3.5 log_10_ copies/mL of viral sgRNA) at 5 and 6 dpe in the nasopharynx for 3 and 2 animals respectively (Figure 6B).

**Figure 6.**
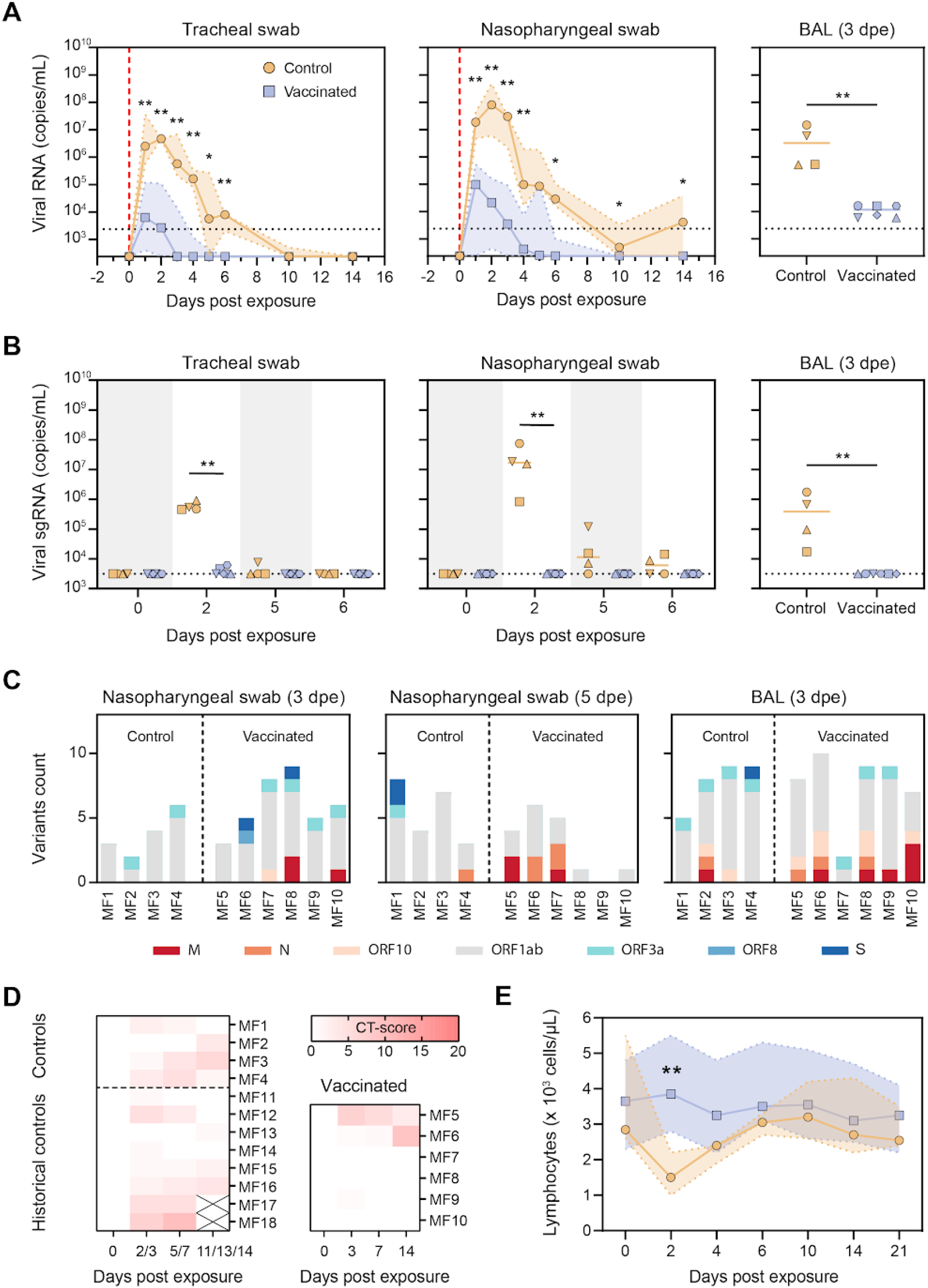
Protective efficacy of SARS-CoV-2 S-I53-50NPs in cynomolgus macaques. (A) Median RNA viral loads in tracheal swabs (left), nasopharyngeal swabs (middle) of control and vaccinated macaques after challenge. The shaded area indicates the range. Viral loads in control and vaccinated macaques after challenge in BAL (right). Bars indicate median viral loads. Vertical red dotted lines indicate the day of challenge. Horizontal dotted lines indicate the limit of quantification. (B) sgRNA viral loads in tracheal swabs (left), nasopharyngeal swabs (middle) and BAL (right) of control and vaccinated macaques after challenge. Bars indicate median viral loads. Dotted line indicates the limit of detection. (C) Viral variants found by viral sequencing, in nasopharyngeal swabs at 3 dpe (left) and 5 dpe (middle), and in BAL at 3 dpe. Colors indicate the ORFs in which mutations were found, as depicted in the legend below. For a list of all identified variants see table S2. (D) Lung CT-scores of control and vaccinated macaques over the course of 14 dpe. CT score includes lesion type (scored from 0 to 3) and lesion volume (scored from 0 to 4) summed for each lobe. (E) Median lymphocyte counts over time in the blood of control and vaccinated macaques after challenge. Shaded area indicates the range. Symbols are the same as indicated in the left panel in A. (A,B,E) Groups were compared using the Mann-Whitney U test (*, *p* < 0.05; **, *p* < 0.01).

In vaccinated macaques, median peak viral loads were 300-fold lower in tracheal samples (6.8 log_10_ vs 4.3 log_10_; *p*= 0.0095) and 500-fold lower in nasopharyngeal samples compared to unvaccinated controls (7.9 log_10_ vs 5.2 log_10_; *p*= 0.0095). With the exception of MF7, all vaccinated animals had undetectable loads at 6 dpe in tracheal and nasopharyngeal swabs (Figure 6A and Figure S4). In addition to the upper airways, SARS-CoV-2 S-I53-50NP vaccination significantly decreased viral loads in the lower airways, as demonstrated by a 275-fold lower median viral load in the bronchoalveolar lavage (BAL) (6.5 log_10_ vs 4.1 log_10_; *p*= 0.0095). Viral replication was also significantly reduced in vaccinated animals. Only 2 out of 6 vaccinated animals (MF5 and MF6) showed detectable sgRNA in the trachea at 2 dpe and median viral loads were 160-fold lower than in control animals (5.7 log_10_ vs 3.5 log_10_; *p*= 0.0095). In the nasopharynx, sgRNA remained below the limit of detection at any of the timepoints. At 2 dpe, median sgRNA loads in the vaccinated animals were 5400-fold lower than in controls (7.2 log_10_ vs 3.5 log_10_; *p*= 0.0048) (Figure 6B). In BAL samples we observed a 120-fold reduction of median sgRNA at day 3 dpe (5.6 log_10_ vs 3.5 log_10_; *p*= 0.0048) (Figures 6A and 6B).

An anamnestic response after challenge (i.e. an increase in (N)Ab titers following challenge after vaccination) implies that vaccination is unable to induce sterilizing immunity. We observed no increase in NAb titers in vaccinated macaques at 2, 3 and 6 weeks after challenge, in contrast to the control animals (Figure S5). Instead, NAbs titers continued to wane, suggesting that vaccine-induced immunity rapidly controlled infection following challenge, preventing a boost of the immune system.

To assess the potential emergence of viral escape mutants in macaques after challenge, viral variants in the challenge inoculum, nasal swabs at 3 and 5 dpe, and in BAL at 3 dpe were sequenced. Two main viral variants were identified in the inoculum (V367F in the S protein and G251V in ORF3a), which were later also found in the nasopharyngeal swabs and BAL samples. A median of 6 subclonal mutations were found per sample over all corresponding time points and anatomical sites, but no major differences were observed between control and vaccinated animals (Figure 6C). The majority of the variants observed in ORF1ab were mostly missense mutations, and consisted of a C>T nucleotide change (Figure S6 and Table S2). A distinct variant in the S sequence arose in 2 vaccinated macaques at day 3 in the nasopharyngeal swab but disappeared at day 5 post challenge, suggesting that no NAb escape mutations emerged in vaccinated animals (Figure 6C).

### Vaccinated cynomolgus macaques have reduced clinical manifestations

Similar to previous observations (Maisonnasse et al., 2020; Yu et al., 2020), during the first 14 dpe all contemporaneous and historical control animals showed mild pulmonary lesions characterized by non-extended ground-glass opacities (GGOs) detected by chest computed tomography (CT) (Figure 6D). By contrast, only 3 out of 6 vaccinated animals showed low CT scores characteristics of mild and non-extended GGOs. Of note, the vaccinated macaque (MF6) with the highest CT score at day 14 showed the lowest S protein- and RBD-specific binding titers, the lowest NAb titers, and the highest viral load and sgRNA at day 5 pe. Whereas all control animals experienced lymphopenia at 2 dpe, corresponding probably to the installation of the response to infection, lymphocyte counts remained stable after challenge for the vaccinated macaques (Figure 6E and Figure S5), in agreement with the absence of detectable anamnestic response. Together, these data further support that vaccination with SARS-CoV-2 S-I53-50NPs reduces the severity of infection.

## Discussion

The development and distribution of a protective vaccine is paramount to bring the SARS-CoV-2 pandemic to a halt. Over the last months, numerous vaccine candidates of different modalities have entered clinical and preclinical studies, including inactivated virus, DNA, mRNA, vector and protein-based vaccines (Klasse et al., 2020). Multivalent presentation of RSV and influenza antigens on two-component self-assembling protein nanoparticles has generated remarkably potent immune responses in non-human primates (Boyoglu-Barnum et al., 2020; Marcandalli et al., 2019). Recently, I53-50NPs presenting the RBD of SARS-CoV-2 induced potent NAb titers and significantly decreased viral loads in humanized mice (Walls et al., 2020b). Here, we show that I53-50NPs presenting twenty copies of pre-fusion SARS-CoV-2 S protein induce robust NAb responses in mice, rabbits and cynomolgus macaques. Vaccination of the latter prevented lymphopenia and significantly reduced viral loads and replication in both the upper and lower respiratory tract, suggesting that SARS-CoV-2 S-I53-50NPs could reduce the risk of severe SARS-CoV-2-associated pathology in vaccinated humans and control viral shedding and transmission.

Evidence is mounting that NAb titers are the immunological correlate of protection for SARS-CoV-2 (Addetia et al., 2020; Yu et al., 2020), and it is increasingly accepted that a successful SARS-CoV-2 vaccine will need to induce potent NAb responses. We observed notable differences in NAb titers by our vaccine and previously described candidates, although comparisons may be biased by, differences in vaccination schedules, assay variability and inconsistencies in data presentation. Here, SARS-CoV-2 S-I53-50NP vaccinated macaques neutralized authentic virus with a median ID_50_ of approximately 4,000 after the final immunization, while NAb titers induced by adenovirus vector vaccines (ChAdOx1 (vanDoremalen et al., 2020) and Janssen (Mercado et al., 2020)), DNA vaccines (Yu *et al.* (Yu et al., 2020) and Inovio (Patel et al., 2020)), and inactivated vaccines (Sinopharm (Wang et al.,2020) and Sinovacc (Gao et al., 2020)) were at least 10-fold lower. However, NAb titers induced by SARS-CoV-2 S-I53-50NPs were similar to Moderna’s mRNA vaccine (Corbett et al., 2020) and lower than Novavax’ and Clover Biopharmaceuticals’ protein vaccines (Corbett et al., 2020; Guebre-Xabier et al., 2020; Liang et al., 2020).

Besides protecting an individual from COVID-19, a key component of an effective SARS-CoV-2 vaccine will be its ability to prevent viral transmission. Hence, sgRNA levels in the upper and lower airways are valuable endpoints in the evaluation of vaccine candidates. Similar to the Janssen and Novavax vaccines, vaccination with SARS-CoV-2 S-I53-50NPs decreased median sgRNA to undetectable levels in the upper airways of all vaccinated animals within 5 dpe, while several other vaccines were unsuccessful in achieving such an effect (vanDoremalen et al., 2020; Patel et al., 2020; Yu et al., 2020). One should consider that the Janssen and Novavax studies used 10- to 100-fold lower challenge doses than used here. Even though three immunizations were used in our vaccine regimen, the high NAb titers at week 6 indicate that two immunizations may be sufficient to confer protection. Collectively, the rapid decrease in sgRNA levels, low level of viral mutations in the S protein, and the absence of an anamnestic response after challenge, emphasize SARS-CoV-2 S-I53-50NP vaccination’s profound ability to control infection and replication.

We propose two factors that may have been responsible for the potent humoral responses and protective efficacy by SARS-CoV-2 S-I53-50NPs. First, the high density of antigen on the I53-50NPs may have facilitated efficient activation of relevant NAb B cell lineages, which is in line with previous studies (Antanasijevic et al., 2020; Brouwer et al., 2019), and is supported by B cell activation experiments described here. Second, by using an S protein that has been stabilized in the prefusion state, we have likely improved the conformation of key NAb epitopes (such as the RBD) and decreased the exposure of non-NAb epitopes. Indeed, it has recently been shown that introduction of the two appropriately positioned prolines and removal of the polybasic-cleavage site significantly improved the protective ability of an S protein vaccine (Amanat et al., 2020). Nonetheless, these mutations alone might not be sufficient to generate stable trimeric S proteins and introducing additional stabilizing mutations, such as the previously described HexaPro mutations (Hsieh et al., 2020), may further improve the SARS-CoV-2 S-I53-50NP-induced humoral responses. Recently, nasal immunization has been shown to dramatically improve SARS-CoV-2 vaccine efficacy over intramuscular dosing (Hassan et al., 2020). Using this alternative administration route may allow SARS-CoV-2 S-I53-50NPs to elicit protective NAb titers in the mucosa and could advance its protective efficacy towards fully sterilizing immunity.

## Acknowledgements

We thank B. Delache, S. Langlois, J. Demilly, N. Dhooge, P. Le Calvez, M. Potier, F. Relouzat, J. M. Robert, T. Prot and C. Dodan for the NHP experiments; L. Bossevot, M. Leonec, L. Moenne-Loccoz and J. Morin for the RT-qPCR and ELISpot assays, and for the preparation of reagents; B. Fert for her help with the CT scans; M. Barendji, J. Dinh and E. Guyon for the NHP sample processing; S. Keyser for the transports organization; N. Dimant and B. Targat for their help with the experimental studies in the context of COVID-19-induced constraints; F. Ducancel and Y. Gorin for their help with the logistics and safety management; I. Mangeot for here help with resources management; The following reagent was obtained through the NIH AIDS Reagent Program, Division of AIDS, NIAID, NIH; Ramos B cells from Drs. L. Wu and V. N. KewalRaman. We thank A. McGuire for kindly sharing the pRRL.EuB29 lentiviral vector to transduce Ramos B cells. We thank P. Bieniasz for kindly sharing the pHIV-1_NL43_ΔENV-NanoLuc and SARS-CoV-2-S_Δ19_ plasmids. We thank H. Nijhuis for sample transportation. We thank A. Chung for sharing knowledge on the Luminex assay protocol and B. Wines and M. Hogarth for sharing the FcγR dimers. We thank Antoine Nougairede for sharing the plasmid used for the sgRNA assays standardization. Finally we thank Dietmar Katinger and Philipp Mundsperger for providing the squalene emulsion and MPLA liposome adjuvants. Animal images in figures 3 and 4 were created with BioRender.com.

## Author contributions

P.J.M.B., M.B., P.M.: Conceptualization, Methodology, Validation, Formal analysis, Investigation, Data curation, Writing - Original Draft, Visualization, Project administration; N.D.B.: Methodology, Investigation, Formal analysis, Supervision, Validation, Writing - Review & Editing M.G., M. C., M.d.G., J.D.A., Y.W.: Conceptualization, Methodology, Investigation, Writing - Original Draft; R.M., V.Ch., S.D.: Methodology, Investigation, Formal analysis, Supervision, Writing - Review & Editing; T.N.: Formal analysis, Writing - Original Draft; J.L.: Investigation, Formal analysis, Writing - Review & Editing; G.K., J.M.G., H.T., A.Z.M.: Investigation, Writing - Original Draft; N.K., K.v.d.S., C.A.v.d.L., Y.A., I.B., J.A.B., M.P., E.E.S., M.J.v.B., T.G.C., J.v.S., N.M.A.O., R.R.: Investigation; K.S.: Conceptualization, Writing - Original Draft; J.V.: Methodology; Y.v.d.V., M.J.v.G.: Conceptualization, Methodology, Supervision, Project administration; G.J.d.B.: Resources, Project administration; V.Co.: Supervision; C.C., R. H.T.F., B.L.H., N.P.K., M.C., A.B.W.: Recourses, Supervision, Writing - Review & Editing; S. v.d.W., E.G.: Resources, Supervision; R.L.G, R.W.S.: Conceptualization, Validation, Resources, Writing - Review & Editing, Supervision, Project administration, Funding acquisition.

## Data and materials availability

The data supporting the findings of the study are available from the corresponding authors upon reasonable request.

## Materials and Methods

### Construct design

To create the SARS-CoV-2 S-I53-50A.1NT1 construct, the previously described pPPI4 plasmid encoding the prefusion SARS-CoV-2 S protein (Brouwer et al., 2020) was digested with PstI and BamHI and ligated in a PstI-BamHI-digested pPPI4 plasmid encoding a modified I53-50A.1NT1 sequence. The original I53-50A.1NT1 plasmid was described previously (Brouwer et al., 2019). Modifications constitute the introduction of GSLEHHHHHH after the final residue to introduce a C-terminal histidine-tag. For ELISAs, Luminex assays, and ELISpots histidine-tagged versions of SARS-CoV-2 S protein and RBD were used, as previously described (Brouwer et al., 2020). To generate S proteins for FACS analyses, the previously described pPPI4 plasmid, encoding a prefusion S ectodomain of SARS-CoV-2 followed by an Avi- and histidine tag, was used. The RBD probe was generated by digesting this plasmid with PstI-BamHI and introducing, by Gibson assembly, a gene encoding the RBD (residues 319-541) directly downstream of a TPA leader sequence.

### Protein expression and purification

All constructs were expressed by transient transfection of HEK 293F cells (Invitrogen) maintained in Freestyle medium (Life Technologies) at a density of 0.8-1.2 million cells/mL. On the day of transfection, a mix of PEImax (1 μg/μL) with expression plasmids (312.5 μg/L) in a 3:1 ratio in OptiMEM (Gibco) were added to the cells. Six days post transfection, supernatants were centrifuged for 30 min at 4000 rpm, filtered using 0.22 μm Steritop filters (Merck Millipore), and subjected to affinity purification using Ni-NTA agarose beads. Protein eluates were then concentrated and buffer exchanged to PBS using Vivaspin filters with the appropriate molecular weight cutoff (GE Healthcare). Protein concentrations were determined by the Nanodrop method using the proteins peptidic molecular weight and extinction coefficient as determined by the online ExPASy software (ProtParam).

### SARS-CoV-2 S-I53-50NP assembly

Ni-NTA eluates of SARS-CoV-2 S-I53-50A.1NT1 (expressed as described above) were buffer exchanged to Tris-buffer-saline (TBS), sterile filtered (0.22 μm) and applied to a Superose 6 increase 10/300 GL column (GE healthcare) in TBS supplemented with 5% glycerol. Size exclusion fractions between 12-14 mL were pooled and equimolar amounts of I53-50B.4PT1, expressed and purified as described elsewhere (Walls et al., 2020b), was added. After an overnight incubation at 4°C, the assembly mix was applied to a Superose 6 increase 10/300 GL column in TBS+5% glycerol to remove unassembled components. Fractions corresponding to assembled SARS-CoV-2 S-I53-50NPs (8.5-9.5 mL) were collected and concentrated using Vivaspin columns with a molecular weight cutoff of 10,000 kDa (GE Healthcare). Protein concentrations were determined by the Nanodrop method using the proteins peptidic molecular weight and extinction coefficient as determined by the online ExPASy software (ProtParam).

### BN-PAGE analysis

BN-PAGE was performed as described previously (de Taeye et al., 2015). Briefly, 2.5 μg of S protein was mixed with loading dye and run on a 4-12% Bis-Tris NuPAGE gel (Invitrogen).

### Negative-stain EM

SARS-CoV-2 S-153-50NPs were added to carbon-covered 400 mesh copper grids and stained with 2% uranyl formate. Micrographs were imaged on a Tecnai F12 Spirit microscope with a 4k FEI Eagle CCD. Leginon (Potter et al., 1999) and Appion (Lander et al., 2009) were used to collect and process micrographs.

### BLI assay

SARS-CoV-2 S-I53-50A.1NT1 and SARS-CoV-2 S-I53-50NP were diluted to 100 nM and 5nM, respectively, in BLI running buffer (PBS/0.1% bovine serum albumin/0.02% Tween20) and antibody binding was assessed using a ForteBio Octet K2. The binding assays were performed at 30°C and with agitation set at 1000 rpm. Antibody was loaded on protein A sensors (ForteBio) at 10 μg/mL in running buffer until a binding threshold of 1 nm was reached. Association and dissociation were measured for 300 s.

### Glycopeptide analysis by liquid chromatography-mass spectrometry

Gycopeptide analysis was essentially performed as described previously (Watanabe et al.,2020). A 30 μg aliquot of SARS-CoV-2 S-I53-50A.1NT1 was denatured, alkylated, reduced and cleaved by the proteases trypsin, chymotrypsin and alpha-lytic protease. Clean-up of the resulting peptide/glycopeptides was performed using C18 Zip-tips and the sample was analysed by nanoLC-ESI MS with an Easy-nLC 1200 (Thermo Fisher Scientific) system coupled to a Fusion mass spectrometer (Thermo Fisher Scientific) using higher energy collision-induced dissociation (HCD) fragmentation. Peptides were separated using an EasySpray PepMap RSLC C18 column (75 μm × 75 cm). Glycopeptide fragmentation data were extracted from the raw file using Byonic™ (Version 3.5) and Byologic™ software (Version 3.5; Protein Metrics Inc.).

### Generation of B cells that stably express COVID-specific B cell receptors

The B cell specific expression plasmid was constructed by exchanging the gl2-1261 gene of the pRRL EuB29 gl2-1261 IgGTM.BCR.GFP.WPRE plasmid (McGuire et al., 2014) with heavy and light chain genes of either COVA1-18 and COVA2-15 using Gibson assembly (Integrated DNA Technologies). The production of lentivirus in HEK 293T and the subsequent transduction was conducted as described elsewhere (ter Brake et al., 2006). In short, lentiviruses were produced by co-transfecting the expression plasmid with pMDL, pVSV-g and pRSV-Rev into HEK 293T cells using lipofectamine 2000 (Invitrogen). Two days post transfection, IgM-negative Ramos B cells cultured in RPMI (Gibco) supplemented with 10% fetal calf serum, streptomycin (100 μg/mL) and penicillin (100 U/mL) (RPMI++) were transduced with filtered (0.45 μm) and concentrated (100 kDa molecular weight cutoff, GE Healthcare) HEK 293T supernatant. Seven days post transduction, BCR-expressing B cells were FACS sorted on IgG and GFP double-positivity using a FACS Aria-II SORP (BD biosciences). B cells were then expanded and cultured indefinitely.

### B cell activation assay

B cell activation experiments of SARS-CoV-2 S protein-specific Ramos B cells were performed as previously described (Brouwer et al., 2019; Sliepen et al., 2019). In short, 4 million cells/mL in RPMI++ were loaded with 1.5 μM of the calcium indicator Indo-1 (Invitrogen) for 30 min at 37°C, washed with Hank’s Balance Salt Solution supplemented with 2 mM CaCl2, followed by another incubation of 30 min at 37°C. Antigen-induced Ca^2+^ influx of COVID-specific B cells were monitored on a LSR Fortessa (BD Biosciences) by measuring the 379/450 nm emission ratio of Indo-1 fluorescence upon UV excitation. Following 30 s of baseline measurement, aliquots of 1 million cells/mL were then stimulated for 100 s at RT with either 5 μg/mL, 1 μg/mL, 200 ng/mL or 40 ng/mL of SARS-CoV-2 S-I53-50A.1NT1 or the equimolar amount presented on I53-50NPs. Ionomycin (Invitrogen) was added to a final concentration of 1 mg/mL to determine the maximum Indo-1-fluorescence. Kinetics analyses were performed using FlowJo v10.7.

### Ethics and biosafety statement animal studies

Cynomolgus macaques (*Macaca fascicularis*), aged 56-66 months and originating from Mauritian AAALAC certified breeding centers were used in this study. All animals were housed in IDMIT infrastructure facilities (CEA, Fontenay-aux-roses), under BSL-2 and BSL-3 containment when necessary (Animal facility authorization #D92-032-02, Préfecture des Hauts de Seine, France) and in compliance with European Directive 2010/63/EU, the French regulations and the Standards for Human Care and Use of Laboratory Animals, of the Office for Laboratory Animal Welfare (OLAW, assurance number #A5826-01, US). The protocols were approved by the institutional ethical committee “Comité d’Ethique en Expérimentation Animale du Commissariat à l’Energie Atomique et aux Energies Alternatives” (CEtEA #44) under statement number A20-011. The study was authorized by the “Research, Innovation and Education Ministry” under registration number APAFIS#24434-2020030216532863v1.

Female New Zealand White rabbits of 2.5-3 kg from multiple litters were used. Animals were sourced and housed at Covance Research Products, Inc. (Denver, PA, USA) and immunizations were performed under permits with approval number C0084-20. Immunization procedures complied with all relevant ethical regulations and protocols of the Covance Institutional Animal Care and Use Committee.

Female Balb/c mice were housed at the Animal Research Institute Amsterdam under BSL-2 conditions. All experiments were performed in accordance with the Dutch Experiment on Animals Act and were approved by the Animal Ethics Committee of the Amsterdam UMC (Permit number 17-4045).

### Animals and study designs

Cynomolgus macaques were randomly assigned in two experimental groups. The vaccinated group (n=6) received 50 ug of SARS-CoV-2 S-I53-50NP adjuvanted with 500 μg of MPLA liposomes (Polymun Scientific, Klosterneuburg, Austria) diluted in PBS at weeks 0, 4 and 10, while control animals (n=4) received no vaccination. Vaccinated animals were sampled in blood, nasal swabs and saliva at weeks 0, 2, 4, 6, 8, 10 and 12. At week 12, all animals were exposed to a total dose of 10^6^ pfu of SARS-CoV-2 virus via the combination of intranasal and intra-tracheal routes (day 0), using atropine (0.04 mg/kg) for pre-medication and ketamine (5 mg/kg) with medetomidine (0.042 mg/kg) for anesthesia. Saliva, as well as nasopharyngeal, tracheal and rectal swabs, were collected at days 1, 2, 3, 4, 5, 6, 10, 14 and 21 days past exposure (dpe) while blood was taken at days 2, 4, 6, 10, 14 and 21 dpe. Bronchoalveolar lavages were performed using 50 mL sterile saline on 3 dpe. Chest CT was performed at baseline and on 3, 7 and 14 dpe in anesthetized animals using tiletamine (4 mg/kg) and zolazepam (4 mg/kg). Blood cell counts, haemoglobin and haematocrit were determined from EDTA blood using a HMX A/L analyzer (Beckman Coulter).

Female New Zealand White rabbits were given two intramuscular immunizations, one in each quadricep, at weeks 0, 4, and 12. The immunization mixture involved 39 μg of SARS-CoV-2-S-I53-50NPs (equal to 30 μg of SARS-CoV-2-S) formulated 1:1 in Squalene Emulsion adjuvant (Polymun, Klosterneuburg, Austria). The rabbits were bled on the day of immunization and then at weeks 2, 4, 6, and 14.

Twelve week old female Balb/c mice received subcutaneous immunizations into the neck skinfold at weeks 0, 4, and 12. The immunization mixture contained 13 μg of SARS-CoV-2-S-I53-50NPs (equal to 10 μg of SARS-CoV-2-S) adjuvanted with 50 μg of polyinosinic-polycytidylic acid (Poly-IC; Invivogen). Blood was collected at weeks-1,2, 6 and 14. Two out of eight mice were sacrificed at week 6 to allow interim analysis of the induced humoral response.

### Patient samples

Sera were collected through the COVID-19 Specific Antibodies (COSCA) study, which was performed as described previously at the Amsterdam University Medical Centre, location AMC, the Netherlands under approval of the local ethical committee of the AMC (NL 73281.018.20) (Brouwer et al., 2020). Briefly, patients with at least one nasopharyngeal swab positive for SARS-CoV-2 as determined by qRT-PCR (Roche LightCycler480, targeting the Envelope-gene 113bp) were included. A venipuncture was performed to obtain blood for serum collection approximately four weeks after onset of COVID-19 symptoms.

### ELISAs

Ni-NTA plates (Qiagen) were loaded with 2 μg/mL of SARS-CoV-2 S protein in TBS for 2 h at RT. Next, washed plates were blocked for 1 h with TBS/2% skimmed milk. Four-fold serial dilutions of inactivated sera, starting from a 1:200 dilution for mice and rabbit sera, and 1:100 dilution for macaque sera, were added in TBS/2% skimmed milk/20% sheep serum and incubated for 2 h at RT. Following plate washing, a 1:3000 dilution of secondary antibody in TBS/2% skimmed milk was added for 1 h at RT. For mice, rabbit and macaque sera horseradish peroxidase (HRP)-labeled goat anti-mice IgG, HRP-labeled goat anti-rabbit IgG and HRP-labeled goat anti-human IgG (Jackson Immunoresearch) was used, respectively. Finally, after washing plates with TBS/0.05% Tween-20, developing solution (1% 3,3’,5,5’-tetranethylbenzidine (Sigma-Aldrich), 0.01% H2O2, 100 mM sodium acetate and 100 mM citric acid) was added to each well and a colorimetric endpoint was obtained by adding 0.8 M H2SO4 after 1.5 min. Binding endpoint titers were determined using a cutoff of 5x background.

As affinity differences between species-specific secondary Abs may bias comparisons between endpoint titers of macaque and convalescent human sera, we used a different ELISA setup to compare S protein-and RBD-specific binding responses. Specifically, ELISA binding responses are compared to a standard curve of species-specific polyclonal IgG so that a semi-quantitative measure of specific IgG concentrations can be obtained. High-binding plates were direct-coated overnight at 4°C with 2 μg/mL of SARS-CoV-2 S protein or 0.4 μg/mL of SARS-CoV-2 RBD in PBS. For capture of the standard curve (i.e. macaque and human polyclonal IgG), plates were coated with 1:2000 (for capture of polyclonal macaque IgG) or 1:1000 (for capture of polyclonal human IgG) of goat anti-Human IgG λ and κ (Southern Biotech). The next day, plates were washed with TBS than contained 0.05% Tween-20 (TBST) and blocked for 1 h with Casein buffer (ThermoScientific). A 1:100, 1:1000 and 1:10,000 dilution of macaque or human serum in Casein buffer were then added to the wells containing S protein or RBD. Five-fold serial dilutions of polyclonal macaque (Molecular Innovations) or human IgG, starting at a concentration of 1 μg/mL, were added to wells containing the coated goat anti-Human IgG λ and κ. Standards and samples were applied in duplicate. Following a 1 h incubation at RT, the plates were washed with TBST and secondary antibody diluted in Casein buffer was added for 1 h. For the plates containing human sera and standard, a 1:3000 dilution of HRP-labeled goat anti-human IgG (Jackson Immunoresearch) was used. For plates containing macaque sera and standard, a 1:50,000 dilution of biotinylated mouse anti-monkey IgG was used (Southern Biotech), followed by a 1 h incubation with a 1:3000 dilution of HRP-labeled streptavidin (Biolegend) in Casein. Finally, after washing plates with TBST, developing solution (1% 3,3’,5,5’-tetranethylbenzidine (Sigma-Aldrich), 0.01% H2O2, 100 mM sodium acetate and 100 mM citric acid) was added to each well and a colorimetric endpoint was obtained by adding 0.8 M H2SO4 after 5 min. OD450-values that fell within the linear range of the standard curve were fitted and the concentration of SARS-CoV-2 S protein- and RBD-specific IgG was determined.

### Pseudovirus neutralization assay

Neutralization assays and the generation of a SARS-CoV-2 pseudovirus containing a NanoLuc luciferase reporter gene were performed as described elsewhere (Schmidt et al.,2020). Briefly, HEK 293T cells (ATCC, CRL-11268) were transfected with a pHIV-1_NL43_ΔENV-NanoLuc reporter virus plasmid and a SARS-CoV-2-SΔ19 plasmid. Cell supernatant containing the pseudoviruses was harvested 48 h post transfection, centrifuged for 5 min at 500 x *g* and sterile filtered through a 0.22 μm pore size PVDF syringe filter. For neutralization assays HEK 293T expressing the SARS-CoV-2 receptor ACE2 (HEK 293T/ACE2) were cultured in DMEM (Gibco), supplemented with 10% fetal bovine serum (FBS), penicillin (100 U/mL), and streptomycin (100 μg/mL).To determine the neutralization activity in serum, saliva or nasopharyngeal samples, HEK 293T/ACE2 cells were first seeded in 96-well plates coated with 50 μg/mL poly-l-lysine at a density of 2×10^4^/well in the culture medium described above but with GlutaMax (Gibco) added. The next day, duplicate serial dilutions of heat inactivated samples were prepared in the same medium as used for seeding of cells and mixed 1:1 with pseudovirus. This mixture was incubated at 37°C for 1 h before adding it to the HEK 293T/ACE2 cells in a 1:1 ratio with the cell culture medium. After 48 h, the cells were lysed and luciferase activity was measured in the lysates using the Nano-Glo Luciferase Assay System (Promega). Relative luminescence units (RLU) were normalized to those from cells infected with SARS-CoV-2 pseudovirus in the absence of sera/saliva/swabs. Neutralization titers (ID_50_-values) were determined as the serum dilution at which infectivity was inhibited by 50%.

### Authentic virus neutralization assay

Serum samples were tested for their neutralization activity against SARS-CoV-2 (German isolate; GISAID ID EPI_ISL 406862; European Virus Archive Global #026V-03883) as described previously (Okba et al., 2020). In short, serum samples were serially diluted in 50 μL Opti-MEM I (Gibco), supplemented with GlutaMAX (Gibco) and 100 U/mL penicillin. Serum dilutions were mixed with 400 plaque-forming units of virus to a total of 100 μL. This mixture was incubated at 37°C for 1 h. After incubation, the mixtures were put on Vero-E6 cells (ATCC CRL-1586) and incubated for 1 h. Cells were then washed, Opti-MEM I supplemented with GlutaMAX (Gibco) was added, and incubated for 8 h. Cells were fixed with 4% formaldehyde in PBS and stained with polyclonal rabbit anti-SARS-CoV Ab (Sino Biological) and secondary Alexa488 conjugated goat-anti-rabbit Ab (Invitrogen). Plates were scanned on the Amersham™ Typhoon™ Biomolecular Imager (channel Cy2; resolution 10 μm; GE Healthcare). Data was analyzed using ImageQuant TL 8.2 image analysis software (GE Healthcare).

### Protein coupling to Luminex beads

Proteins were covalently coupled to Magplex beads (Luminex Corporation) using a two-step carbodiimide reaction and a ratio of 75 μg protein SARS-CoV-2 S protein to 12,5 million beads. Magplex beads (Luminex Corporation) were washed with 100 mM monobasic sodium phosphate pH 6.2, activated for 30 min on a rotor at RT by addition of Sulfo-N-Hydroxysulfosuccinimide (Thermo Fisher Scientific) and 1-Ethyl-3-(3-dimethylaminopropyl) carbodiimide (Thermo Fisher Scientific). The activated beads were washed three times with 50 mM MES pH 5.0 and added to SARS-CoV-2 S protein which was diluted in 50 mM MES pH 5.0. The beads and protein were incubated for 3 h on a rotator at RT. Afterwards, the beads were washed with PBS and blocked with PBS containing 0.1% BSA, 0.02% Tween-20 and 0.05% Sodium Azide at pH 7.0 for 30 min on a rotator at RT. Finally, the beads were washed and stored in PBS containing 0.05% Sodium Azide at 4°C and used within 3 months.

### Luminex assays

50 μL of a working bead mixture containing 20 beads per μL was incubated overnight at 4°C with 50 μL of diluted serum, saliva or nasopharyngeal swab. Pilot experiments determined the optimal dilution for detection of SARS-CoV-2 S protein-specific IgG, IgA and IgM antibodies in serum to be 1:50,000, for the detection of antibody binding to FcyRIIa, FcyRIIIa and C1q in serum at 1:500 and for detection of S protein-specific IgG and IgA in saliva and nose fluid at 1:20. Plates were sealed and incubated on a plate shaker overnight at 4°C. The next day, plates were washed with TBS containing 0.05% Tween-20 (TBST) using a hand-held magnetic separator. Beads were resuspended in 50 μL of Goat-anti-human IgG-PE, Goat-anti human IgA-PE or Mouse-anti human IgM-PE (Southern Biotech) and incubated on a plate shaker at RT for 2 h. For C1q binding, beads were resuspended in 50 μL purified human C1q (Complement Technology, Inc.), which was biotinylated and conjugated to Streptavidin-PE. For the detection of FcyRIIa and FcyRIIIa binding, beads were resuspended in 50 μL biotinylated human FcyRIIa and FcyRIIIa ectodomain dimers (courtesy of Bruce Wines and Mark Hogarth) for 2 h incubation, after which the beads were washed with TBST and then incubated with 50 μL Streptavidin-PE (Invitrogen) on a plate shaker for 1 h. Afterwards, the beads were washed with TBST and resuspended in 110 μL Bioplex sheath fluid (Bio-Rad). The beads were agitated for a few minutes on a plate shaker at RT and then read-out was performed on the Bioplex 200 (Bio-Rad). Resulting mean fluorescence intensity (MFI) values were corrected by subtraction of MFI values from buffer and beads only wells. Relative MFI values were obtained by dividing the MFI by the background MFI at week 0. Reproducibility of the results was confirmed by performing replicate runs.

### SARS-CoV-2 S protein-specific Tfh cell analysis using an activation induced marker (AIM) assay

The AIM assay was performed as previously described (Reiss et al., 2017). Cryopreserved macaque PBMCs from 2 weeks after final immunization (week 12) were thawed and counted. 1×10^6^ cells per condition were plated in 96 U-shape well plates (CELLSTAR, Kremsmünster) and rested for 2-3 h in RPMI (Gibco), supplemented with 10% FBS. Subsequently, cells were incubated with 0.5 mg/mL anti-CD40 (Miltenyi Biotec) for 15 min at 37°C, to prevent downregulation of CD40 ligand (CD40L) by blocking CD40-CD40L interaction. PBMCs were stimulated for 18 h at 37°C with 5 mg/mL SARS-CoV-2 S protein. As a negative control no stimulants were added, and as a positive control 1 mg/mL Staphylococcal enterotoxin B was used. Thereafter, PBMCs were washed using FACS buffer (PBS, supplemented with 1% FBS). PBMCs were then incubated with the Tfh-specific chemokine receptor antibody CXCR5 PE-Cy7 (MU5UBEE; eBioscience) for 10 min at 37°C. Next, the following antibodies and viability dye were incubated with PBMCs at RT for 20 min: anti-CD3 BUV395 (clone SP34-2; BD Biosciences), anti-CD4 BUV495 (clone SK3; BD Biosciences), anti-CD154 BV421 (clone TRAP1; BD Biosciences), anti-CD69 BV785 (clone FN50; BD Biosciences), and LiveDead-eF780 (eBioscience). Cells were washed twice with FACS buffer before acquisition on the BD Fortessa II flow cytometer (BD biosciences). Analysis was performed on FlowJo v.10.7.1.

### SARS-CoV-2 S protein- and RBD-specific B cell analysis

Biotinylated SARS-CoV-2 S protein and RBD were conjugated to streptavidin bound fluorophores. Briefly, the biotinylated proteins were incubated for a minimum of 1 h at 4°C with the streptavidin-conjugates AF647 (Biolegend), BV421 (Biolegend) and BB515 (BD Biosciences) at a 1:2 protein to fluorochrome ratio. The fluorescent probes were incubated for at least 10 min with 10mM biotin (GeneCopoeia) to saturate the unconjugated streptavidin-fluorochrome complexes. Cryopreserved macaque PBMCs from 2 weeks after final immunization (week 12) were thawed and counted. 5×10^6^ cells were stained for 30 min at 4°C with the fluorescent probes, a viability marker (LiveDead-eF780, eBiosciences) and the following B cell-specific antibodies: anti-CD20-PE-CF594 (2H7; BD Biosciences), anti-IgG-PE-Cy7 (G18-145;BD Biosciences), anti-CD27-PE (M-T271; BD Biosciences), and anti-IgM-BV605 (MHM-88; Biolegend). Cells were washed twice with FACS buffer and acquired on the FACS-ARIA-SORP 4 laser (BD Biosciences). Analysis was performed on FlowJo v.10.7.1.

### Viruses and cells

For the cynomolgus macaque challenge study, SARS-CoV-2 virus (hCoV-19/France/ lDF0372/2020 strain) was isolated by the National Reference Center for Respiratory Viruses (Institut Pasteur, Paris, France) as described recently (Lescure et al., 2020). Virus stocks used *in vivo* were produced by two passages on Vero E6 cells in DMEM (Gibco) without FBS, supplemented with penicillin at 100 U/mL, streptomycin at 100 μg/mL, and 1 μg/mL TPCK-trypsin at 37°C in a humidified CO2 incubator and titrated on Vero E6 cells.

### Virus quantification in cynomolgus macaque samples

Upper respiratory (nasopharyngeal and tracheal) and rectal specimens were collected with swabs (Viral Transport Medium, CDC, DSR-052-01). Tracheal swabs were performed by insertion of the swab above the tip of the epiglottis into the upper trachea at approximately 1.5 cm of the epiglottis. All specimens were stored between 2°C and 8°C until analysis by RT-qPCR with a plasmid standard concentration range containing an RdRp gene fragment including the RdRp-IP4 RT-PCR target sequence. SARS-CoV-2 E gene subgenomic mRNA (sgmRNA) levels were assessed by RT-qPCR using primers and probes previously described (Corman et al., 2020; Wölfel et al., 2020): leader-specific primer sgLeadSARSCoV2-F CGATCTCTTGTAGATCTGTTCTC, E-Sarbeco-R primer ATATTGCAGCAGTACGCACACA and E-Sarbeco probe HEX-ACACTAGCCATCCTTACTGCGCTTCG-BHQ1. The protocol describing the procedure for the detection of SARS-CoV-2 is available on the WHO website (https://www.who.int/docs/default-source/coronaviruse/real-time-rt-pcr-assays-for-the-detection-of-sars-cov-2-institut-pasteur-paris.pdf?sfvrsn=3662fcb6_2).

### Chest computed tomography and image analysis

Acquisition was done using a computed tomography (CT) system (Vereos-Ingenuity, Philips) as previously described (Maisonnasse et al., 2020).Lesions were defined as ground glass opacity, crazy-paving pattern, consolidation or pleural thickening as previously described (Pan et al., 2020; Shi et al., 2020). Lesions and scoring were assessed in each lung lobe blindly and independently by two persons and final results were made by consensus. Overall CT scores include the lesion type (scored from 0 to 3) and lesion volume (scored from 0 to 4) summed for each lobe as previously described (Maisonnasse et al., 2020).

### ELISpot assays

IFNγ ELISpot assay of PBMCs was performed using the Monkey IFNγ ELISpot PRO kit (Mabtech Monkey IFNγ ELISPOT pro, #3421M-2APT) according to the manufacturer’s instructions. PBMCs were plated at a concentration of 200,000 cells per well and were stimulated with SARS-CoV-2 S protein at a final concentration of 5 μg/mL. Plates were incubated for 42 h at 37°C in an atmosphere containing 5% CO2, then washed 5 times with PBS and incubated for 2 h at 37°C with a biotinylated anti-IFNγ antibody. After 5 washes, spots were developed by adding 0.45 μm-filtered ready-to-use BCIP/NBT-plus substrate solution and counted with an automated ELISpot reader ELRIFL04 (Autoimmun Diagnostika GmbH, Strassberg, Germany). Spot forming units (SFU) per 1.0×10^6^ PBMCs are means of duplicates for each animal.

### Viral sequencing

30 RNA samples from nasal swabs and bronchoalveolar lavage at day 3 and 5 were selected for sequencing. cDNA and multiplex PCR reactions were prepared following the ARTIC SARS-CoV-2 sequencing protocol v2 (Tyson et al., 2020). The inoculum was also sequenced. V3 primer scheme (https://github.com/artic-network/primer-schemes/tree/master/nCoV-2019/V3) was used to perform the multiplex PCR for SARS-CoV-2. All samples were run for 35 cycles in the two multiplex PCRs. Pooled and cleaned PCR reactions were quantified using QubitTM fluorometer (Invitrogen). The Ligation Sequencing kit (SQK-LSK109; Oxford Nanopore Technologies) was used to prepare the library following the manufacturer’s protocol (“PCR tiling of COVID-19 virus”, release F; Oxford Nanopore Technologies). 24 samples were multiplexed using Native Barcoding Expansion 1-12 and Native Barcoding Expansion 13-24 kits (EXP-NBD104 and EXP-NBD114; Oxford Nanopore Technologies). Two libraries of 24 samples were prepared independently and quantified by QubitTM fluorometer (Invitrogen). After the quality control, two R9.4 flowcells (FLO-MIN106; Oxford Nanopore Technologies) were primed as described in the manufacturer’s protocol and loaded with 45 and 32 ng of library. Sequencing was performed on a GridION (Oxford Nanopore Technologies) for 72 h with high-accuracy Guppy basecalling (v3.2.10). After sequencing, demultiplexing was performed using Guppy v4.0.14 with the option--require_barcodes_both_ends to ensure high quality demultiplexing. Reads were then filtered by Nanoplot v1.28.1 based on length and quality to select high quality reads. Then, reads were aligned on the SARS-CoV-2 reference genome NC_045512.2 using minimap2 v2.17. Primary alignments were filtered based on reads length alignment and reads identity. Finally, variant calling was performed with Longshot v0.4.1. Identified variants were then analyzed using in-house scripts. Variants absent from the inoculum but present in one or more samples were selected for further analysis.

### Statistical analysis

Binding titers (endpoint titers and concentrations) and midpoint neutralization titers were determined using Graphpad Prism 7.0. Comparisons between two experimental groups were made using a Mann-Whitney U test (Excel 2016, Graphpad Prism 7.0).

---

**Figure S1.**
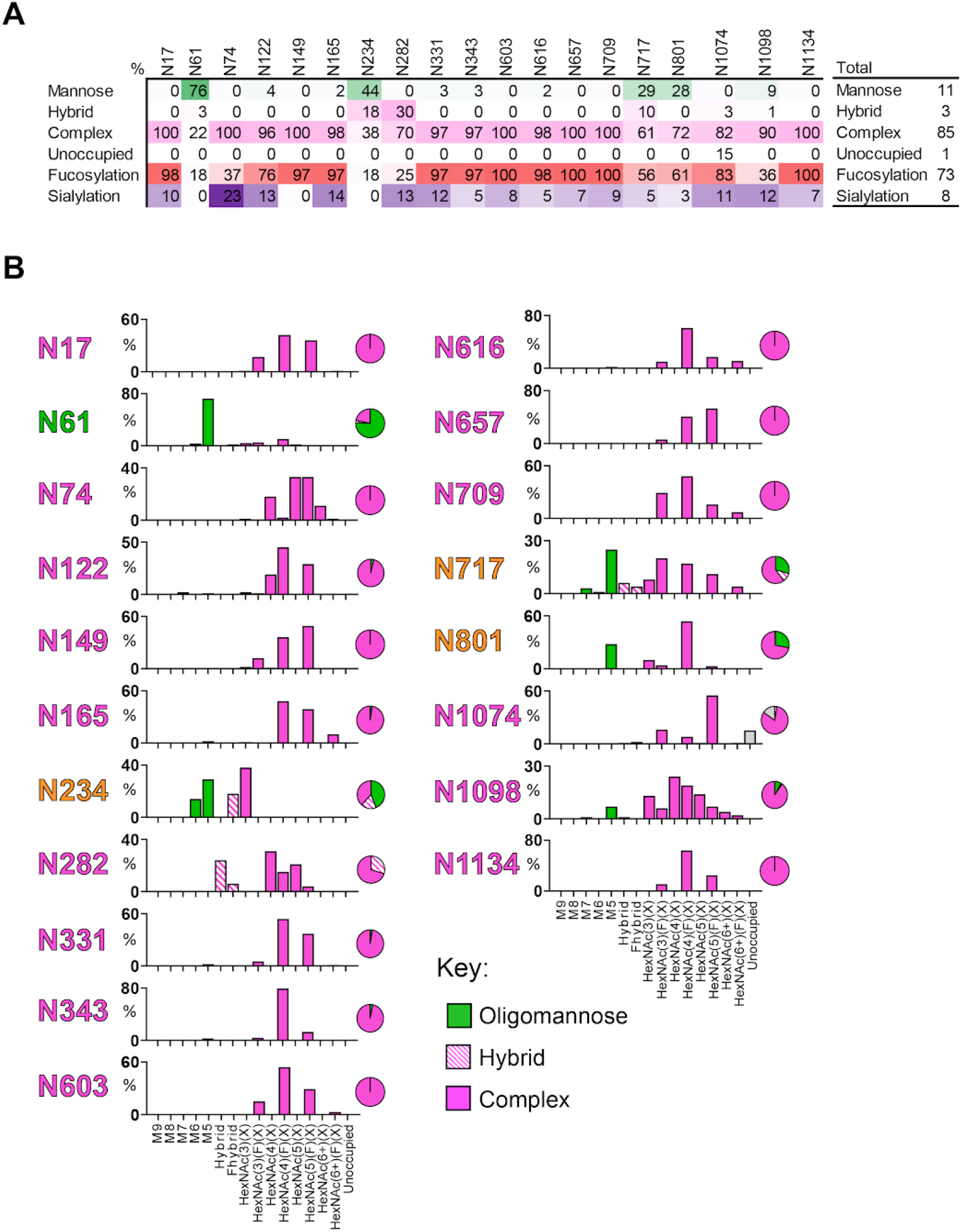
Site-specific glycan analysis of SARS-CoV-2 S I53-50NPs, related to Figure 1. (A) The table categorizes the glycan compositions into oligomannose-, hybrid-, and complex-type as well as the percentage of glycan species containing at least one fucose or one sialic acid residue. The overall averages are shown in the right-hand table. (B) Site-specific distribution of N-linked glycans. The graphs summarize quantitative mass spectrometric analysis of the glycan population present at individual N-linked glycosylation sites simplified into categories of glycans. The oligomannose-type glycan series (M9 to M5; Man9GlcNAc2 to Man5GlcNAc2) is colored green, afucosylated and fucosylated hybrid-type glycans (Hybrid & F Hybrid) dashed pink, and complex glycans grouped according to the number of antennae and presence of core fucosylation (A1 to FA4) and are colored pink. Unoccupancy of an N-linked glycan site is represented in grey. The pie charts summarize the quantification of these glycans. Glycan sites are colored according to oligomannose-type glycan content with the glycan sites labelled in green (80-100% oligomannose), orange (30-79% oligomannose) and pink (0-29% oligomannose).

**Figure S2.**
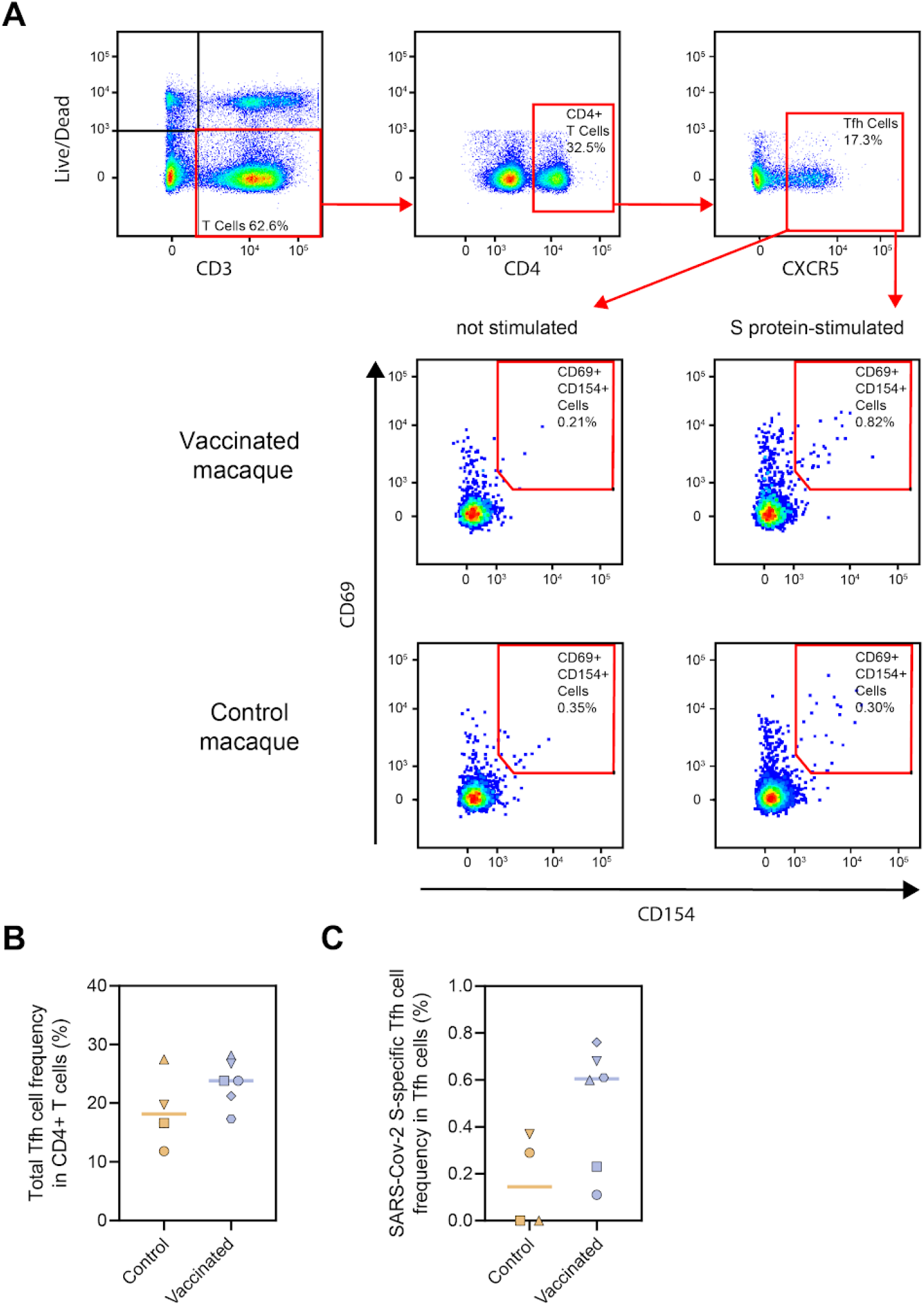
SARS-CoV-2 S protein-specific Tfh cell responses in control and vaccinated macaques, related to Figure 4. (A) Representative gating strategy for the identification of SARS-CoV-2 S protein-specific Tfh cells. PBMCs were stimulated overnight with SARS-CoV-2 S protein and Tfh activation was assessed the next day by analyzing CD69 and CD154 expression. (B) Frequency of total Tfh cells in CD4+ T cell population following stimulation with SARS-CoV-2 S protein. (C) Frequency of SARS-CoV-2 S-protein specific Tfh cells within the total Tfh cell population. Corresponding background (i.e. frequency of activated Tfh cells in non-stimulated cells) has been subtracted from each data point.

**Figure S3.**
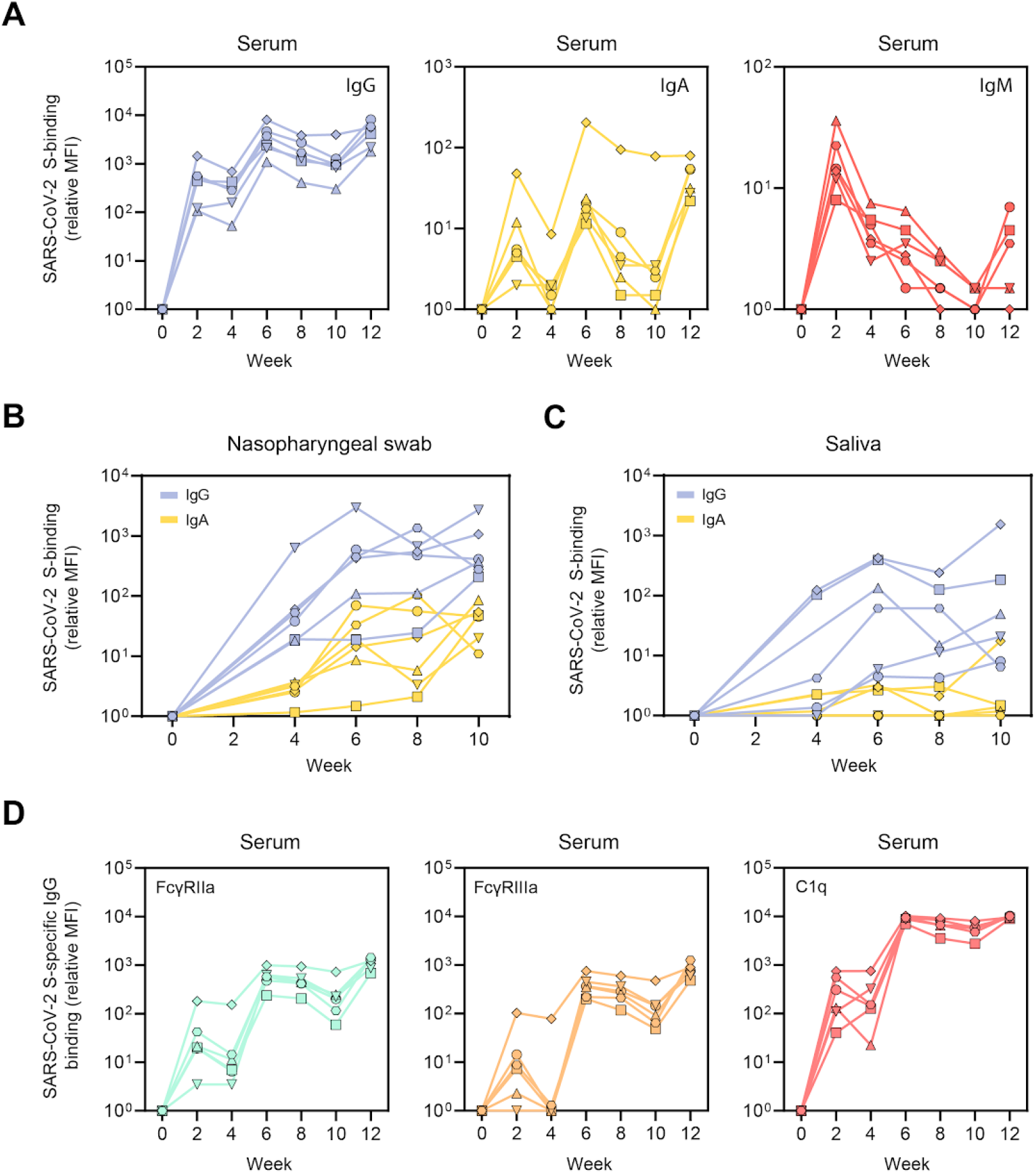
SARS-CoV-2 S protein-specific Ig levels and Fc-receptor binding in vaccinated cynomolgus macaques in samples from diverse anatomical sites, related to Figure 5. (A) Relative MFI of IgG (left), IgA (middle) and IgM (right) binding to SARS-CoV-2 S protein measured with a Luminex-based serology assay in serum samples. (B) Relative MFI of IgG and IgA binding to SARS-CoV-2 S protein in nasopharyngeal swabs. (C) Relative MFI of IgG and IgA binding to SARS-CoV-2 S protein in saliva. (D) Relative MFI of FcγRIIa (left), FcγRIIIa (middle) and C1q (right) binding to SARS-CoV-2 S protein-specific IgG in serum samples.

**Figure S4.**
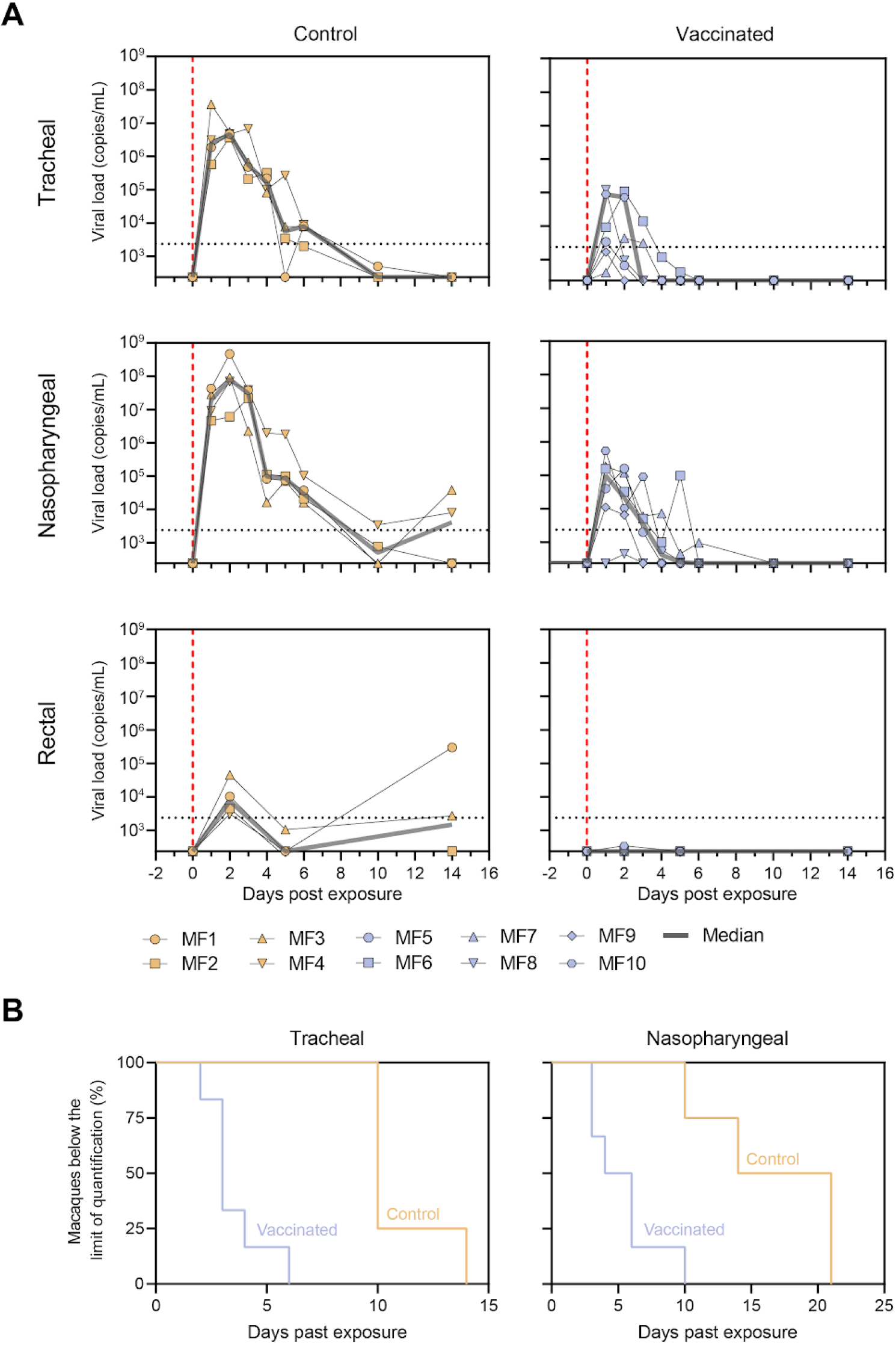
Viral loads in control and vaccinated cynomolgus macaques after SARS-CoV-2 challenge, related to Figure 6. (A) RNA viral loads in tracheal swabs (top), nasopharyngeal swabs (middle) and rectal samples (bottom) of control (left) and SARS-CoV-2 S-I53-50NP vaccinated macaques (right) after challenge. The grey line represents the median viral load. Vertical red dotted lines indicate the day of challenge. Horizontal dotted lines indicate the limit of quantification. Symbols identify individual macaques. (B) Percentage of macaques in which the RNA viral loads drop below the limit of quantification over time in tracheal swabs (left) and nasopharyngeal swabs (right).

**Figure S5.**
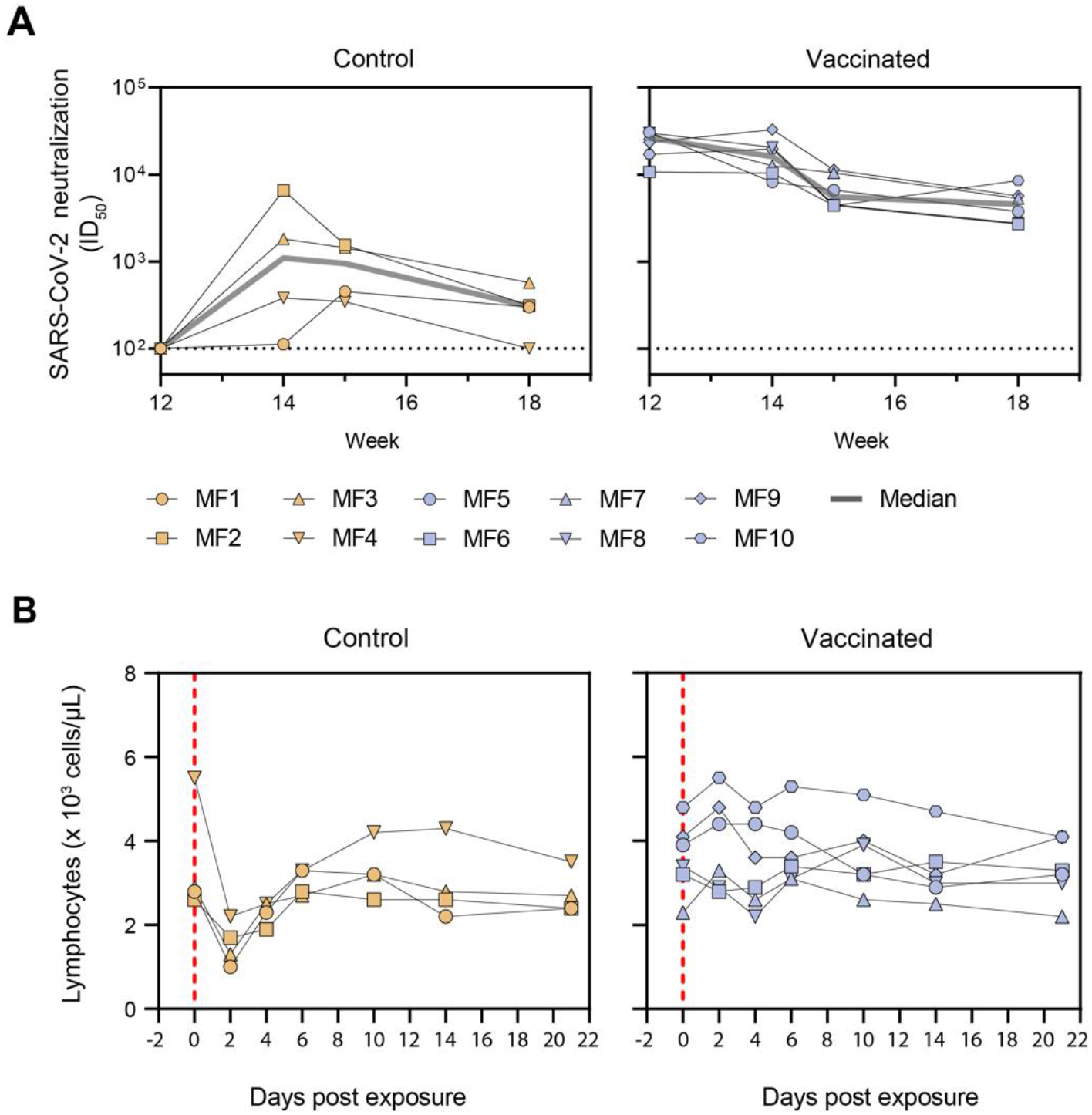
Anamnestic immune response and lymphocyte counts in control and vaccinated cynomolgus macaques after SARS-CoV-2 challenge, related to Figure 6. (A) SARS-CoV-2 pseudovirus neutralization titers. The grey line represents the median titers over-time. Symbols identify individual macaques. (B) Lymphocyte counts over time in the blood of control and SARS-CoV-2 S-I53-50NP vaccinated macaques after challenge. Vertical red dotted lines indicate the day of challenge. Symbols identify individual macaques, as indicated in panel A.

**Figure S6.**
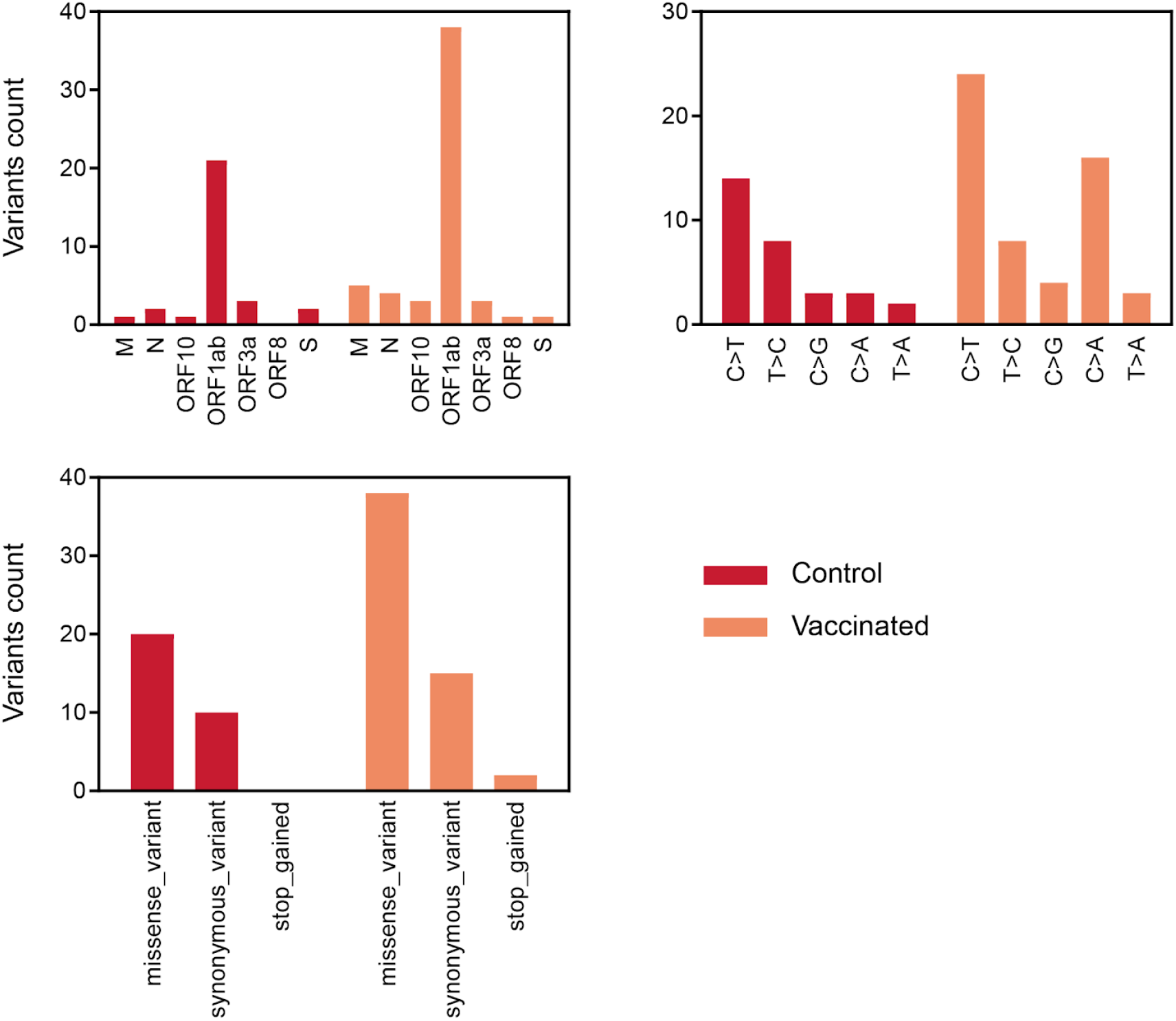
Viral sequence variants found in control and vaccinated cynomolgus macaques after SARS-CoV-2 challenge, related to Figure 6. Sum of viral variants found by viral sequencing, in nasopharyngeal swabs at 3 dpe and 5 dpe, and in BAL at 3 dpe, specified for the ORF in which it was found (top left) the type of nucleotide change (top right) and the effect is has on the amino acid sequence (bottom; missense_variant = amino acid change, synonymous_variant = no amino acid change, stop_gained = introduction of a stop codon). For a list of all identified variants see table S3.

**Table S1. SARS-CoV-2 pseudovirus and authentic virus neutralization titers (ID_50_s) in mice and rabbits, related to figure 3.**

Auxiliary Supplementary Material

**Table S2. SARS-CoV-2 pseudovirus and authentic virus neutralization titers (ID_50_s) in cynomolgus macaques, related to figure 5.**

Auxiliary Supplementary Material

**Table S3. Viral variants identified by viral sequencing in control and vaccinated cynomolgus macaques after challenge, related to figure 6.**

Auxiliary Supplementary Material

