## Supplementary material for "Two-component spike nanoparticle vaccine protects macaques from SARS-CoV-2 infection": Table S1

Table S1 SARS-CoV-2 pseudovirus and authentic virus serum neutralization ID50s in BLAB/c mice and rabbits

Pseudovirus neutralization

| | Animal ID | Week -1/0 | Week 6 | Week 14 |
|---|---|---|---|---|
| Mice | 1 | <100* | 75063 | n.d. |
| | 2 | | 25593 | 63015 |
| | 3 | <100* | 2588 | 231100 |
| | 4 | | 23399 | 63418 |
| | 5 | <100* | 2992 | n.d. |
| | 6 | | 8179 | 17235 |
| | 7 | <100* | 14035 | 35046 |
| | 8 | | 19548 | 18749 |
| Rabbits | 1 | <100 | 68298 | 34813 |
| | 2 | <100 | 41968 | 76139 |
| | 3 | <100 | 41162 | 135128 |
| | 4 | <100 | 77885 | 152796 |
| | 5 | <100 | 90884 | 186147 |

Authentic virus neutralization

| | Animal ID | Week 14 |
|---|---|---|
| Mice | 1 | n.d. |
| | 2 | 6700 |
| | 3 | 9716 |
| | 4 | 4651 |
| | 5 | n.d. |
| | 6 | 1649 |
| | 7 | 3478 |
| | 8 | 3068 |
| Rabbits | 1 | 4514 |
| | 2 | 28180 |
| | 3 | 15110 |
| | 4 | 10138 |
| | 5 | 30851 |

| | |
|---|---|
| n.d. | Not determined |
| | <100 |
| | 101 - 1,000 ID50 |
| | 1,001 - 10,000 ID50 |
| | 10,001 - 100,000 ID50 |
| | >100,001 ID50 |

*Samples were pooled
