## Supplementary material for "Two-component spike nanoparticle vaccine protects macaques from SARS-CoV-2 infection": Table S2

Table S2 SARS-CoV-2 pseudovirus and authentic virus serum neutralization ID50s in cynomolgus macaques

Pseudovirus neutralization

| | Animal ID | Week 0 | Week 2 | Week 4 | Week 6 | Week 8 | Week 10 | Week 12 | Week 14 | Week 15 | Week 18 |
|---|---|---|---|---|---|---|---|---|---|---|---|
| Control macaques | MF1 | n.d. | n.d. | n.d. | n.d. | n.d. | n.d. | <100 | 112 | 452 | 299 |
| | MF2 | n.d. | n.d. | n.d. | n.d. | n.d. | n.d. | <100 | 6583 | 1553 | 311 |
| | MF3 | n.d. | n.d. | n.d. | n.d. | n.d. | n.d. | <100 | 1818 | 1442 | 571 |
| | MF4 | n.d. | n.d. | n.d. | n.d. | n.d. | n.d. | <100 | 381 | 344 | <100 |
| Vaccinated macaques | MF5 | <100 | 162 | 251 | 11340 | 9179 | 3043 | 30766 | 8212 | 6615 | 3782 |
| | MF6 | <100 | 912 | <100 | 2981 | 1324 | 1469 | 10720 | 10384 | 4415 | 2690 |
| | MF7 | <100 | 584 | <100 | 10171 | 6646 | 2780 | 29198 | 12679 | 10476 | 5322 |
| | MF8 | <100 | 599 | <100 | 5087 | 3894 | 3150 | 30080 | 20526 | 4507 | 2762 |
| | MF9 | <100 | 183 | <100 | 22450 | 30300 | 22328 | 23523 | 32909 | 11352 | 5680 |
| | MF10 | <100 | <100 | 2515 | 7558 | 2991 | 1101 | 17128 | 19730 | 4376 | 8544 |

Authentic virus neutralization

| | Animal ID | Week 0 | Week 6 | Week 12 |
|---|---|---|---|---|
| Control macaques | MF1 | n.d. | n.d. | n.d. |
| | MF2 | n.d. | n.d. | n.d. |
| | MF3 | n.d. | n.d. | n.d. |
| | MF4 | n.d. | n.d. | n.d. |
| Vaccinated macaques | MF5 | <20 | 792 | 4190 |
| | MF6 | <20 | 166 | 1239 |
| | MF7 | <20 | 2155 | 5095 |
| | MF8 | <20 | 1629 | 1913 |
| | MF9 | <20 | 3040 | 3694 |
| | MF10 | <20 | 1372 | 4851 |

| | |
|---|---|
| n.d. | Not determined |
| | <100 |
| | 101 - 1,000 ID50 |
| | 1,001 - 10,000 ID50 |
| | 10,001 - 100,000 ID50 |
| | >100,001 ID50 |
