## Supplementary material for "Two-component spike nanoparticle vaccine protects macaques from SARS-CoV-2 infection": Table S3

Table S3 All viral variants found by viral sequencing

| sample | time | sample site | sample group | variant position | variant frequency | variant type | annotation | impact |
|---|---|---|---|---|---|---|---|---|
| MF5 | 3 dpe | nasal swab | vaccinated | 18317 | 10% | C>T | ORF1ab:Gly6018Ser | missense_variant |
| MF5 | 3 dpe | nasal swab | vaccinated | 17200 | 5% | T>A | ORF1ab:His5645Gln | missense_variant |
| MF5 | 3 dpe | nasal swab | vaccinated | 494 | 22% | C>T | ORF1ab:Arg77* | stop_gained |
| MF5 | 3 dpe | BAL | vaccinated | 29461 | 50% | T>A | N:Pro396Pro | synonymous_variant |
| MF5 | 3 dpe | BAL | vaccinated | 6473 | 5% | C>A | ORF1ab:Glu2070* | stop_gained |
| MF5 | 3 dpe | BAL | vaccinated | 20557 | 7% | C>G | ORF1ab:Leu6764Leu | synonymous_variant |
| MF5 | 3 dpe | BAL | vaccinated | 5736 | 6% | C>T | ORF1ab:Ala1824Val | missense_variant |
| MF5 | 3 dpe | BAL | vaccinated | 6337 | 11% | C>A | ORF1ab:Ser2024Ser | synonymous_variant |
| MF5 | 3 dpe | BAL | vaccinated | 29613 | 6% | C>A | ORF10:Cys19Phe | missense_variant |
| MF5 | 3 dpe | BAL | vaccinated | 1121 | 20% | C>G | ORF1ab:Pro286Ala | missense_variant |
| MF5 | 3 dpe | BAL | vaccinated | 12917 | 33% | C>A | ORF1ab:Asp4218Tyr | missense_variant |
| MF5 | 5 dpe | nasal swab | vaccinated | 27103 | 14% | C>T | M:Ala194Val | missense_variant |
| MF5 | 5 dpe | nasal swab | vaccinated | 27013 | 14% | T>C | M:Leu164Pro | missense_variant |
| MF5 | 5 dpe | nasal swab | vaccinated | 16768 | 50% | C>T | ORF1ab:Thr5501Thr | synonymous_variant |
| MF5 | 5 dpe | nasal swab | vaccinated | 454 | 10% | T>C | ORF1ab:Gln63Gln | synonymous_variant |
| MF6 | 3 dpe | nasal swab | vaccinated | 17200 | 6% | T>A | ORF1ab:His5645Gln | missense_variant |
| MF6 | 3 dpe | nasal swab | vaccinated | 24096 | 12% | C>T | S:Ala845Val | missense_variant |
| MF6 | 3 dpe | nasal swab | vaccinated | 5595 | 8% | T>C | ORF1ab:Glu1777Gly | missense_variant |
| MF6 | 3 dpe | nasal swab | vaccinated | 8043 | 13% | C>T | ORF1ab:Ala2593Val | missense_variant |
| MF6 | 3 dpe | nasal swab | vaccinated | 28239 | 25% | C>A | ORF8:Val116Phe | missense_variant |
| MF6 | 3 dpe | BAL | vaccinated | 5814 | 50% | C>A | ORF1ab:Gly1850Val | missense_variant |
| MF6 | 3 dpe | BAL | vaccinated | 19168 | 100% | T>C | ORF1ab:Asn6301Asn | synonymous_variant |
| MF6 | 3 dpe | BAL | vaccinated | 16768 | 7% | C>T | ORF1ab:Thr5501Thr | synonymous_variant |
| MF6 | 3 dpe | BAL | vaccinated | 28313 | 100% | C>T | N:Arg14Cys | missense_variant |
| MF6 | 3 dpe | BAL | vaccinated | 29613 | 8% | C>A | ORF10:Cys19Phe | missense_variant |
| MF6 | 3 dpe | BAL | vaccinated | 27084 | 11% | C>T | M:Ala188Thr | missense_variant |
| MF6 | 3 dpe | BAL | vaccinated | 29564 | 5% | T>C | ORF10:Tyr3His | missense_variant |
| MF6 | 3 dpe | BAL | vaccinated | 8043 | 50% | C>T | ORF1ab:Ala2593Val | missense_variant |
| MF6 | 3 dpe | BAL | vaccinated | 17064 | 6% | C>A | ORF1ab:Ser5600Ile | missense_variant |
| MF6 | 3 dpe | BAL | vaccinated | 17200 | 13% | T>A | ORF1ab:His5645Gln | missense_variant |
| MF6 | 5 dpe | nasal swab | vaccinated | 15103 | 6% | C>T | ORF1ab:Leu4946Leu | synonymous_variant |
| MF6 | 5 dpe | nasal swab | vaccinated | 13476 | 6% | C>T | ORF1ab:Ala4404Val | missense_variant |
| MF6 | 5 dpe | nasal swab | vaccinated | 16220 | 7% | C>T | ORF1ab:Arg5319Cys | missense_variant |
| MF6 | 5 dpe | nasal swab | vaccinated | 9309 | 10% | C>T | ORF1ab:Pro3015Leu | missense_variant |
| MF6 | 5 dpe | nasal swab | vaccinated | 29348 | 8% | C>G | N:Ala359Pro | missense_variant |
| MF6 | 5 dpe | nasal swab | vaccinated | 29461 | 9% | T>A | N:Pro396Pro | synonymous_variant |
| MF7 | 3 dpe | nasal swab | vaccinated | 29613 | 5% | C>A | ORF10:Cys19Phe | missense_variant |
| MF7 | 3 dpe | nasal swab | vaccinated | 1296 | 25% | C>A | ORF1ab:Cys344Phe | missense_variant |
| MF7 | 3 dpe | nasal swab | vaccinated | 18079 | 13% | C>A | ORF1ab:Arg5938Ser | missense_variant |
| MF7 | 3 dpe | nasal swab | vaccinated | 5736 | 14% | C>T | ORF1ab:Ala1824Val | missense_variant |
| MF7 | 3 dpe | nasal swab | vaccinated | 9539 | 13% | C>T | ORF1ab:Leu3092Phe | missense_variant |
| MF7 | 3 dpe | nasal swab | vaccinated | 26065 | 20% | C>A | ORF3a:Val225Phe | missense_variant |
| MF7 | 3 dpe | nasal swab | vaccinated | 15327 | 10% | C>A | ORF1ab:Cys5021Phe | missense_variant |
| MF7 | 3 dpe | nasal swab | vaccinated | 15391 | 20% | C>T | ORF1ab:Thr5042Thr | synonymous_variant |
| MF7 | 3 dpe | BAL | vaccinated | 25429 | 50% | C>A | ORF3a:Val13Leu | missense_variant |
| MF7 | 3 dpe | BAL | vaccinated | 17254 | 10% | C>A | ORF1ab:Val5663Val | synonymous_variant |
| MF7 | 5 dpe | nasal swab | vaccinated | 14881 | 25% | C>A | ORF1ab:Leu4872Phe | missense_variant |
| MF7 | 5 dpe | nasal swab | vaccinated | 27084 | 11% | C>T | M:Ala188Thr | missense_variant |
| MF7 | 5 dpe | nasal swab | vaccinated | 29348 | 10% | C>G | N:Ala359Pro | missense_variant |
| MF7 | 5 dpe | nasal swab | vaccinated | 11150 | 100% | C>T | ORF1ab:Val3629Ile | missense_variant |
| MF7 | 5 dpe | nasal swab | vaccinated | 29026 | 8% | T>C | N:Ala251Ala | synonymous_variant |
| MF8 | 3 dpe | nasal swab | vaccinated | 17064 | 7% | C>A | ORF1ab:Ser5600Ile | missense_variant |
| MF8 | 3 dpe | nasal swab | vaccinated | 1121 | 20% | C>G | ORF1ab:Pro286Ala | missense_variant |
| MF8 | 3 dpe | nasal swab | vaccinated | 556 | 6% | C>T | ORF1ab:Tyr97Tyr | synonymous_variant |
| MF8 | 3 dpe | nasal swab | vaccinated | 19168 | 25% | T>C | ORF1ab:Asn6301Asn | synonymous_variant |
| MF8 | 3 dpe | nasal swab | vaccinated | 11150 | 5% | C>T | ORF1ab:Val3629Ile | missense_variant |
| MF8 | 3 dpe | nasal swab | vaccinated | 24096 | 13% | C>T | S:Ala845Val | missense_variant |
| MF8 | 3 dpe | nasal swab | vaccinated | 26594 | 8% | T>C | M:Ile24Met | missense_variant |
| MF8 | 3 dpe | nasal swab | vaccinated | 26065 | 9% | C>A | ORF3a:Val225Phe | missense_variant |
| MF8 | 3 dpe | nasal swab | vaccinated | 27013 | 6% | T>C | M:Leu164Pro | missense_variant |
| MF8 | 3 dpe | BAL | vaccinated | 25419 | 20% | T>A | ORF3a:Thr9Thr | synonymous_variant |
| MF8 | 3 dpe | BAL | vaccinated | 29026 | 6% | T>C | N:Ala251Ala | synonymous_variant |
| MF8 | 3 dpe | BAL | vaccinated | 6307 | 17% | C>G | ORF1ab:Trp2014Cys | missense_variant |
| MF8 | 3 dpe | BAL | vaccinated | 18893 | 14% | C>T | ORF1ab:Ala6210Thr | missense_variant |
| MF8 | 3 dpe | BAL | vaccinated | 27013 | 11% | T>C | M:Leu164Pro | missense_variant |
| MF8 | 3 dpe | BAL | vaccinated | 29613 | 10% | C>A | ORF10:Cys19Phe | missense_variant |
| MF8 | 3 dpe | BAL | vaccinated | 14677 | 11% | C>T | ORF1ab:Pro4804Pro | synonymous_variant |
| MF8 | 3 dpe | BAL | vaccinated | 29654 | 10% | C>A | ORF10:Val33Phe | missense_variant |
| MF8 | 3 dpe | BAL | vaccinated | 16768 | 100% | C>T | ORF1ab:Thr5501Thr | synonymous_variant |
| MF8 | 5 dpe | nasal swab | vaccinated | 15391 | 100% | C>T | ORF1ab:Thr5042Thr | synonymous_variant |
| MF1 | 3 dpe | nasal swab | control | 17200 | 6% | T>A | ORF1ab:His5645Gln | missense_variant |
| MF1 | 3 dpe | nasal swab | control | 18317 | 6% | C>T | ORF1ab:Gly6018Ser | missense_variant |
| MF1 | 3 dpe | nasal swab | control | 13476 | 5% | C>T | ORF1ab:Ala4404Val | missense_variant |
| MF1 | 3 dpe | BAL | control | 556 | 7% | C>T | ORF1ab:Tyr97Tyr | synonymous_variant |
| MF1 | 3 dpe | BAL | control | 25876 | 7% | T>C | ORF3a:Ser162Gly | missense_variant |
| MF1 | 3 dpe | BAL | control | 8830 | 7% | T>C | ORF1ab:Ala2855Ala | synonymous_variant |
| MF1 | 3 dpe | BAL | control | 7588 | 5% | T>C | ORF1ab:Gly2441Gly | synonymous_variant |
| MF1 | 3 dpe | BAL | control | 6884 | 50% | C>T | ORF1ab:Gly2207Ser | missense_variant |
| MF1 | 5 dpe | nasal swab | control | 17658 | 6% | C>G | ORF1ab:Cys5798Ser | missense_variant |
| MF1 | 5 dpe | nasal swab | control | 26065 | 5% | C>A | ORF3a:Val225Phe | missense_variant |
| MF1 | 5 dpe | nasal swab | control | 23623 | 6% | T>C | S:Val687Val | synonymous_variant |
| MF1 | 5 dpe | nasal swab | control | 13476 | 6% | C>T | ORF1ab:Ala4404Val | missense_variant |
| MF1 | 5 dpe | nasal swab | control | 24096 | 5% | C>T | S:Ala845Val | missense_variant |

| | | | | | | | | |
|---|---|---|---|---|---|---|---|---|
| MF1 | 5 dpe | nasal swab | control | 18317 | 5% | C>T | ORF1ab:Gly6018Ser | missense_variant |
| MF1 | 5 dpe | nasal swab | control | 15103 | 8% | C>T | ORF1ab:Leu4946Leu | synonymous_variant |
| MF1 | 5 dpe | nasal swab | control | 5300 | 5% | C>G | ORF1ab:Ala1679Pro | missense_variant |
| MF2 | 3 dpe | nasal swab | control | 26065 | 5% | C>A | ORF3a:Val225Phe | missense_variant |
| MF2 | 3 dpe | nasal swab | control | 17200 | 6% | T>A | ORF1ab:His5645Gln | missense_variant |
| MF2 | 3 dpe | BAL | control | 29466 | 10% | C>T | N:Ala398Val | missense_variant |
| MF2 | 3 dpe | BAL | control | 4255 | 6% | C>T | ORF1ab:Pro1330Pro | synonymous_variant |
| MF2 | 3 dpe | BAL | control | 25419 | 8% | T>A | ORF3a:Thr9Thr | synonymous_variant |
| MF2 | 3 dpe | BAL | control | 26594 | 6% | T>C | M:Ile24Met | missense_variant |
| MF2 | 3 dpe | BAL | control | 556 | 6% | C>T | ORF1ab:Tyr97Tyr | synonymous_variant |
| MF2 | 3 dpe | BAL | control | 1121 | 5% | C>G | ORF1ab:Pro286Ala | missense_variant |
| MF2 | 3 dpe | BAL | control | 29613 | 6% | C>A | ORF10:Cys19Phe | missense_variant |
| MF2 | 3 dpe | BAL | control | 17658 | 11% | C>G | ORF1ab:Cys5798Ser | missense_variant |
| MF2 | 5 dpe | nasal swab | control | 556 | 5% | C>T | ORF1ab:Tyr97Tyr | synonymous_variant |
| MF2 | 5 dpe | nasal swab | control | 16220 | 5% | C>T | ORF1ab:Arg5319Cys | missense_variant |
| MF2 | 5 dpe | nasal swab | control | 13476 | 6% | C>T | ORF1ab:Ala4404Val | missense_variant |
| MF2 | 5 dpe | nasal swab | control | 15103 | 6% | C>T | ORF1ab:Leu4946Leu | synonymous_variant |
| MF10 | 3 dpe | nasal swab | vaccinated | 13476 | 18% | C>T | ORF1ab:Ala4404Val | missense_variant |
| MF10 | 3 dpe | nasal swab | vaccinated | 556 | 6% | C>T | ORF1ab:Tyr97Tyr | synonymous_variant |
| MF10 | 3 dpe | nasal swab | vaccinated | 18317 | 5% | C>T | ORF1ab:Gly6018Ser | missense_variant |
| MF10 | 3 dpe | nasal swab | vaccinated | 27103 | 20% | C>T | M:Ala194Val | missense_variant |
| MF10 | 3 dpe | nasal swab | vaccinated | 26065 | 8% | C>A | ORF3a:Val225Phe | missense_variant |
| MF10 | 3 dpe | nasal swab | vaccinated | 5814 | 5% | C>A | ORF1ab:Gly1850Val | missense_variant |
| MF10 | 3 dpe | BAL | vaccinated | 6021 | 25% | C>T | ORF1ab:Pro1919Leu | missense_variant |
| MF10 | 3 dpe | BAL | vaccinated | 14479 | 11% | C>T | ORF1ab:Thr4738Thr | synonymous_variant |
| MF10 | 3 dpe | BAL | vaccinated | 29654 | 6% | C>A | ORF10:Val33Phe | missense_variant |
| MF10 | 3 dpe | BAL | vaccinated | 27084 | 20% | C>T | M:Ala188Thr | missense_variant |
| MF10 | 3 dpe | BAL | vaccinated | 8172 | 50% | C>T | ORF1ab:Ala2636Val | missense_variant |
| MF10 | 3 dpe | BAL | vaccinated | 27103 | 17% | C>T | M:Ala194Val | missense_variant |
| MF10 | 3 dpe | BAL | vaccinated | 27076 | 17% | T>C | M:Gln185Arg | missense_variant |
| MF10 | 5 dpe | nasal swab | vaccinated | 14479 | 25% | C>T | ORF1ab:Thr4738Thr | synonymous_variant |
| MF3 | 3 dpe | nasal swab | control | 18317 | 6% | C>T | ORF1ab:Gly6018Ser | missense_variant |
| MF3 | 3 dpe | nasal swab | control | 17200 | 6% | T>A | ORF1ab:His5645Gln | missense_variant |
| MF3 | 3 dpe | nasal swab | control | 2860 | 13% | T>C | ORF1ab:Gly865Gly | synonymous_variant |
| MF3 | 3 dpe | nasal swab | control | 4255 | 5% | C>T | ORF1ab:Pro1330Pro | synonymous_variant |
| MF3 | 3 dpe | BAL | control | 2909 | 8% | T>C | ORF1ab:Thr882Ala | missense_variant |
| MF3 | 3 dpe | BAL | control | 2860 | 15% | T>C | ORF1ab:Gly865Gly | synonymous_variant |
| MF3 | 3 dpe | BAL | control | 1985 | 6% | T>C | ORF1ab:Ser574Pro | missense_variant |
| MF3 | 3 dpe | BAL | control | 18079 | 5% | C>A | ORF1ab:Arg5938Ser | missense_variant |
| MF3 | 3 dpe | BAL | control | 16768 | 5% | C>T | ORF1ab:Thr5501Thr | synonymous_variant |
| MF3 | 3 dpe | BAL | control | 556 | 5% | C>T | ORF1ab:Tyr97Tyr | synonymous_variant |
| MF3 | 3 dpe | BAL | control | 13476 | 8% | C>T | ORF1ab:Ala4404Val | missense_variant |
| MF3 | 3 dpe | BAL | control | 29613 | 6% | C>A | ORF10:Cys19Phe | missense_variant |
| MF3 | 3 dpe | BAL | control | 26065 | 8% | C>A | ORF3a:Val225Phe | missense_variant |
| MF3 | 5 dpe | nasal swab | control | 556 | 7% | C>T | ORF1ab:Tyr97Tyr | synonymous_variant |
| MF3 | 5 dpe | nasal swab | control | 7588 | 6% | T>C | ORF1ab:Gly2441Gly | synonymous_variant |
| MF3 | 5 dpe | nasal swab | control | 2860 | 10% | T>C | ORF1ab:Gly865Gly | synonymous_variant |
| MF3 | 5 dpe | nasal swab | control | 15103 | 5% | C>T | ORF1ab:Leu4946Leu | synonymous_variant |
| MF3 | 5 dpe | nasal swab | control | 17200 | 5% | T>A | ORF1ab:His5645Gln | missense_variant |
| MF3 | 5 dpe | nasal swab | control | 13476 | 6% | C>T | ORF1ab:Ala4404Val | missense_variant |
| MF3 | 5 dpe | nasal swab | control | 1121 | 5% | C>G | ORF1ab:Pro286Ala | missense_variant |
| MF4 | 3 dpe | nasal swab | control | 15103 | 5% | C>T | ORF1ab:Leu4946Leu | synonymous_variant |
| MF4 | 3 dpe | nasal swab | control | 17200 | 6% | T>A | ORF1ab:His5645Gln | missense_variant |
| MF4 | 3 dpe | nasal swab | control | 18317 | 6% | C>T | ORF1ab:Gly6018Ser | missense_variant |
| MF4 | 3 dpe | nasal swab | control | 6884 | 100% | C>T | ORF1ab:Gly2207Ser | missense_variant |
| MF4 | 3 dpe | nasal swab | control | 6078 | 6% | C>T | ORF1ab:Ala1938Val | missense_variant |
| MF4 | 3 dpe | nasal swab | control | 26065 | 5% | C>A | ORF3a:Val225Phe | missense_variant |
| MF4 | 3 dpe | BAL | control | 25876 | 5% | T>C | ORF3a:Ser162Gly | missense_variant |
| MF4 | 3 dpe | BAL | control | 15103 | 9% | C>T | ORF1ab:Leu4946Leu | synonymous_variant |
| MF4 | 3 dpe | BAL | control | 1121 | 6% | C>G | ORF1ab:Pro286Ala | missense_variant |
| MF4 | 3 dpe | BAL | control | 14677 | 6% | C>T | ORF1ab:Pro4804Pro | synonymous_variant |
| MF4 | 3 dpe | BAL | control | 18079 | 5% | C>A | ORF1ab:Arg5938Ser | missense_variant |
| MF4 | 3 dpe | BAL | control | 2860 | 6% | T>C | ORF1ab:Gly865Gly | synonymous_variant |
| MF4 | 3 dpe | BAL | control | 18317 | 6% | C>T | ORF1ab:Gly6018Ser | missense_variant |
| MF4 | 3 dpe | BAL | control | 24096 | 6% | C>T | S:Ala845Val | missense_variant |
| MF4 | 3 dpe | BAL | control | 556 | 5% | C>T | ORF1ab:Tyr97Tyr | synonymous_variant |
| MF4 | 5 dpe | nasal swab | control | 8043 | 25% | C>T | ORF1ab:Ala2593Val | missense_variant |
| MF4 | 5 dpe | nasal swab | control | 18317 | 5% | C>T | ORF1ab:Gly6018Ser | missense_variant |
| MF4 | 5 dpe | nasal swab | control | 28313 | 6% | C>T | N:Arg14Cys | missense_variant |

dpe : days post exposure
missense_variant : mutation that causes change in aminoacid
stop_gained : mutation that introduces stop codon
synonymous_variant : silent mutation
